# Interleukin-10 suppression enhances T-cell antitumor immunity and responses to checkpoint blockade in chronic lymphocytic leukemia

**DOI:** 10.1101/2020.07.15.204560

**Authors:** J.R. Rivas, Y. Liu, S.S. Alhakeem, J.M. Eckenrode, F. Marti, J.P. Collard, Y. Zhang, K.A. Shaaban, N. Muthusamy, G.C. Hildebrandt, R.A. Fleischman, L. Chen, J.S. Thorson, M. Leggas, S. Bondada

## Abstract

T-cell dysfunction is a hallmark of B-cell Chronic Lymphocytic Leukemia (CLL), where CLL cells downregulate T-cell responses through regulatory molecules including programmed death ligand-1 (PD-L1) and Interleukin-10 (IL-10). Immune checkpoint blockade (ICB) aims to restore T-cell function by preventing the ligation of inhibitory receptors like PD-1. However, most CLL patients do not respond well to this therapy. Thus, we investigated whether IL-10 suppression could enhance antitumor T-cell activity and responses to ICB. Since CLL IL-10 expression depends on Sp1, we utilized a novel, better tolerated analogue of the Sp1 inhibitor mithramycin (MTM_ox_32*E*) to suppress CLL IL-10. MTM_ox_32*E* treatment inhibited mouse and human CLL IL-10 production and maintained T-cell effector function *in vitro*. In the Eμ-Tcl1 mouse model, treatment reduced plasma IL-10 and CLL burden and increased CD8^+^ T-cell proliferation, effector and memory cell prevalence, and interferon-γ production. When combined with ICB, suppression of IL-10 improved responses to anti-PD-L1 as shown by a 4.5-fold decrease in CLL cell burden compared to anti-PD-L1 alone. Combination therapy also produced more interferon-γ^+^, cytotoxic effector KLRG1^+^, and memory CD8^+^ T-cells, and fewer exhausted T-cells. Since current therapies for CLL do not target IL-10, this provides a novel strategy to improve immunotherapies.

## Introduction

B-cell Chronic Lymphocytic Leukemia (CLL) is a malignancy of CD5^+^CD19^+^ B-lymphocytes that leads to severe immune dysfunction. T-cells play multiple roles in the CLL microenvironment and disease progression, from aiding CLL growth to lacking functionality which impedes immunity. CLL cells decrease T-cell antitumor immunity by expressing regulatory molecules including checkpoint ligands and anti-inflammatory cytokines like interleukin-10 (IL-10)(1-5). Patient T-cells then proliferate less after stimulation, increase inhibitory receptor expression, and experience impaired immune synapse formation(1, 6, 7). These functional defects skew T-cell subset distributions, alter T-cell metabolism, and create pseudo-exhausted T-cells(1, 8). Ultimately this results in prevalent infections and secondary cancers, causing much of CLL patient morbidity and mortality(9). Since CLL is rarely curable, there is a need for new approaches to restore immunity.

After repeated antigen stimulation, T-cells upregulate checkpoint receptors such as programmed death-1 (PD-1) that decrease T-cell receptor sensitivity. Ligation of these receptors decreases CD8^+^ T-cell antitumor immunity via T-cell exhaustion(1). Antibody-based immune checkpoint blockade (ICB) can increase T-cell activity by preventing this interaction. Pembrolizumab (anti-PD-1) ICB was tested to reestablish T-cell functionality in high risk CLL patients with limited success. Only those with Richter’s transformation responded(10), suggesting additional mechanisms downregulate T-cell antitumor immunity. Even adoptively transferred anti-CD19 chimeric antigen receptor (CAR) T-cells upregulate apoptosis and exhaustion genes in most CLL patients, resulting in surprisingly low response rates(11). Hence several studies are exploring combination therapies to improve responses to T-cell immunotherapy in CLL(12-14).

Since IL-10 is a well-known anti-inflammatory cytokine, we reasoned that CLL-derived IL-10 may suppress host antitumor T-cell responses(3, 4). Elevated IL-10 levels correlate with aggressive disease in CLL and other cancers(15, 16), and CLL patients with elevated plasma IL-10 have a worse 3-year survival rate than those with low levels(15). Furthermore, reduced DNA methylation at the IL-10 locus or polymorphisms that increase IL-10 production are correlated with decreased survival in CLL(17, 18). Despite the importance of IL-10 in the microenvironment and improved antitumor responses with IL-10 blockade in other cancer types(19-21), no small molecules blocking IL-10 are currently available(22, 23). Previously, we showed that IL-10R deletion improved CD8^+^ T-cell-efficacy to control CLL growth in the Eμ-TCL1 mouse CLL model(4). Eμ-TCL1 mice express the oncogene T-cell leukemia 1 (TCL1) under the μ-enhancer and the VH promoter leading to an aggressive CLL-like disease in 9 to 11 months(24). Like their human counterparts, murine CLL cells secrete IL-10 in a BCR dependent manner, which can be interrupted by small molecules that inhibit BCR signaling(3-5).

We hypothesized that reducing IL-10 would restore host anti-tumor responses and improve the efficacy of T-cell-based immunotherapy. We suppressed CLL-derived IL-10 with anti-IL-10 or by inhibiting Sp1, a transcription factor required for CLL IL-10 production(4), with a novel analog of mithramycin (MTM_ox_32*E*, structure in supplementary methods). MTM_ox_32*E* exhibits reduced toxicity and increased potency over the parent molecule in other cancer models(25, 26). This treatment significantly decreased CLL burden in the blood, spleen and bone marrow, and when combined with anti-PD-L1, suppressing IL-10 reduced tumor burden and enhanced T-cell functionality over ICB alone. Thus, suppressing IL-10 could potentially improve antitumor T-cell function in human CLL and other immunosuppressive cancers.

## Methods

### Human samples

Patient blood and healthy donor leukopaks were obtained with informed consent in accordance with the regulations of the Institutional Review Board of the University of Kentucky Research Foundation. Diagnosis of CLL was assigned by board-certified hematologists (Table S1). Peripheral blood mononuclear cells (PBMCs) were isolated by Ficoll-Paque density centrifugation, then used fresh for healthy donor cells and thawed from frozen CLL samples. CD45^+^CD5^+^CD19^+^ CLL cell expression of CD38, Zap70 and CD49d were evaluated by flow cytometry (antibodies in Table S2) on an LSR II flow cytometer (BD Biosciences, Franklin Lakes, NJ). Mutational status of the expressed BCR was determined by PCR from PBMC RNA, as previously described(4).

### Mice

Eμ-TCL1 (C57BL/6J background), NOD.SCID IL-2Rγ^-/-^ (NSG), and C57BL/6J mice were bred in house. Experimental groups were randomized by including 8-12 week-old mice of both genders (numbers in figure legends). All animal studies were approved by the University of Kentucky Institutional Animal Care and Use Committee (IACUC). Three to four million CD19^+^ bead purified Eμ-TCL1 cells were adoptively transferred into NSG mice with or without primed CD8^+^ T-cells at 32:1 (CLL:T-cell). Cell sorts were performed with MojoSort Mouse Pan B Isolation Kit II (BioLegend, San Diego, CA) for CD19+ cells and with Miltenyi’s magnetic-activated cell sorting (MACS) T-cell subset Isolation Kits (Miltenyi Biotec, Bergisch Gladbach, Germany) for CD4+ and CD8+ cells according to the manufacturers’ protocols. To prime CD8^+^ T-cells, ten million Ficoll-Paque purified Eμ-TCL1 splenocytes (>90% CD5^+^CD19^+^) were adoptively transferred into C57BL/6J mice and ten days later T-cells were purified from the spleen. Disease burden was monitored in the blood by CD45^+^CD5^+^CD19^+^ frequency on an LSR II flow cytometer (BD Biosciences) and CD45^+^ cell count on a MACSQuant VYB Flow Cytometer (Miltenyi Biotec). Mice were treated with 12mg/kg MTM_ox_32*E* or vehicle (5% Kolliphor EL, 1% DMSO) by tail vein injection, 150μg/mouse anti-IL-10 (clone JES5-2A5, BioXCell, West Lebanon, NH), or 10mg/kg anti-PD-L1 or isotype control by intraperitoneal injection (clone B7-H1, BioXCell). Investigators were not blinded. When CD5^+^CD19^+^ cells comprised more than 70% of the peripheral blood, mice were euthanized. For *in vivo* proliferation, 1mg of BrdU (Sigma Aldrich, St. Louis, MO) in 100µL PBS was injected intraperitoneally three hours before euthanasia.

### Cell culture

Eμ-TCL1 splenocytes, human CLL PBMCs (hCLL), and sorted T-cell subsets from mice and humans were cultured at 1×10^6^/mL for 24 hours in 96 well plates. Concentrations of inhibitors and stimulants are listed in figure legends. Anti-IgM (Jackson ImmunoResearch, West Grove, PA), anti-CD3, anti-CD28 (BioLegend) and MTM (Enzo Life Sciences, Farmingdale, NY) are commercially sourced, while MTM_ox_32*E* was synthesized in house(26). Secreted cytokines were measured with ELISA MAX Standard Set Mouse or Human kits according to the manufacturer’s protocol (BioLegend). Cellular viability was determined by resuspending cells in cRPMI with 0.1mM resazurin (Sigma Aldrich) I, incubating for four hours at 37°C, and measuring fluorescence (ex:560nm, em:590nm) on a SpectraMax M5 plate reader (Molecular Devices, Sunnyvale, CA)(27).

### Proliferation assays

CD8^+^ cells were sorted as above from primed WT mice, healthy donor leukopaks or hCLL PBMCs. Mouse T-cells were cultured at 0.75×10^6^/mL for three days with soluble 10μg/mL anti-CD3 and/or Ficoll-Paque purified Eμ-TCL1 splenocytes. Human T-cells were cultured at 1×10^6^/mL for five days with plate-bound 10μg/mL anti-CD3 and soluble 1μg/mL anti-CD28. Cells were then pulsed with 1μCi ^3^H-thymidine (Perkin Elmer, Waltham, MA) or 40μM BrdU for four hours at 37°C. Counts from ^3^H-thymidine incorporation were measured with a TopCount NXT Microplate Scintillation & Luminescence Counter (Perkin Elmer) and BrdU incorporation was measured on an LSR II or Symphony PRO flow cytometer (BD Biosciences).

### Statistics

*A priori* power analysis determined the number of mice used in experiments. GraphPad Prism 8 software was used for statistical tests (GraphPad Software, San Diego, CA), as listed in each figure legend. One-way ANOVA was used for three or more groups, two-way ANOVA was used for two or more groups over time points or concentrations, and two-tailed student’s t-test was used between two groups. Variance was comparable between groups in the same test.

Additional information is in the Supplementary Materials and Methods.

## Results

### Blocking CLL-derived IL-10 restores T-cell function

Previously we observed that removing CD8^+^ T-cell IL-10 signaling improves control of CLL growth in the Eμ-TCL1 adoptive transfer model(4). Considering IL-10 is a potent regulator of T-cell antitumor immunity(23), we hypothesized that CLL-derived IL-10 inhibits CLL-primed CD8^+^ T-cell responses to antigenic stimuli. We primed C57BL/6J CD8^+^ T-cells with Eμ-TCL1 splenocytes *in vivo* and tested their responses to re-challenge with CLL splenocytes *ex vivo* with and without anti-IL-10. Anti-CD3 induced T-cell proliferation was not changed by anti-IL-10 antibody in the absence of CLL (Fig. 1A). In CLL-T-cell cocultures (3:1), IL-10 levels were only moderately increased with few CLL present, and anti-IL-10 did not change T-cell proliferation (Fig. 1A, 1B). In contrast, proliferation increased four-fold when IL-10 was blocked in cultures with higher CLL:CD8^+^ ratios (Fig. 1A). This is not explained by CLL cell activity, as blocking IL-10 does not enhance Eμ-TCL1 proliferation under similar culture conditions(4). In CLL:T-cell co-cultures IFN-γ levels were very low, which were substantially increased by neutralizing IL-10 (Fig. 1C).

**Figure 1:**
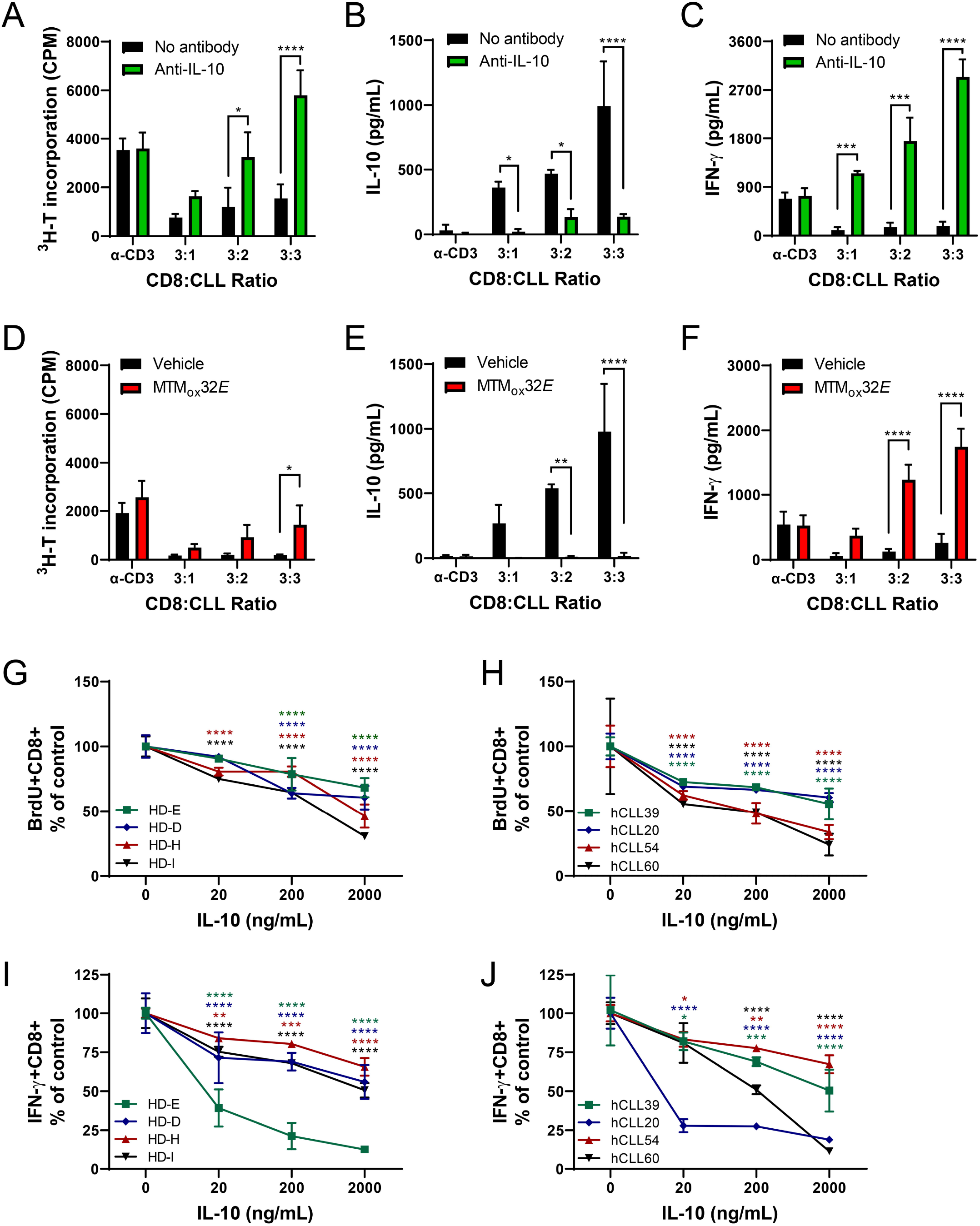
CLL IL-10 suppresses T-cell responses. (A) Proliferation of CLL-primed CD8^+^ T-cells in response to anti-CD3 or varying numbers of Eμ-TCL1 splenocytes after 72 hours of stimulation, with or without 10μg/mL anti-IL-10. Secreted IL-10 (B) and IFN-γ (C) from cultures in panel A (Eμ-TCL1 spleen cells alone produced 986pg/mL IL-10 and 80pg/mL IFN-γ). (D) Proliferation of CLL-primed CD8^+^ T-cells stimulated with anti-CD3 or varying numbers of Eμ-TCL1 splenocytes after 72 hours with or without 1μM MTM_ox_32*E*. Secreted IL-10 (E) and IFN-γ (F) from cultures in panel D (Eμ-TCL1 spleen cells alone produced 948pg/mL IL-10 and 78pg/mL IFN-γ). (G-H) Normalized proliferation (BrdU incorporation) of healthy donor (G) or CLL patient (H) human CD8^+^ T-cells in response to a 5-day stimulation with plate bound anti-CD3 with soluble anti-CD28 and recombinant human IL-10. (I-J) Normalized intracellular IFN-γ production from cells in (G-H), measured by flow cytometry. Frequency of responding cells in G-J were normalized to responses of CD8^+^ T-cells from same donor/patient cultured without IL-10. Two-way ANOVA between control and antibody or inhibitor treatment was used to calculate statistical significance. **p<0.01, ***p<0.001, ****p<0.0001

In earlier studies we showed CLL IL-10 production depends on Sp1 activity(4). Sp1 binds GC rich regions of DNA to regulate transcription, which can be competitively inhibited by MTM or MTM analogues(28). Inhibiting CLL IL-10 with the MTM analogue MTM_ox_32*E* also increased T-cell functionality in our CD8^+^:CLL co-cultures. Proliferation of primed CD8+ T-cells was rescued by MTM_ox_32*E* treatment in the 1:1 co-cultures (Fig. 1D). MTM_ox_32*E* suppressed IL-10 production in all conditions, even in 3:1 co-cultures, where low IL-10 levels only had a small effect on T-cell proliferation (Fig. 1D-E). Drug treatment did not affect anti-CD3 driven proliferation of primed CD8^+^ T-cells in isolation (Fig. 1D). Like co-cultures blocked with anti-IL-10, decreasing IL-10 with MTM_ox_32*E* (Fig. 1E) also increased IFN-γ secretion (Fig. 1F). Notably, 1μM MTM_ox_32*E* treatment did not alter IFN-γ secretion from anti-CD3 stimulated, primed CD8^+^ T-cells in the absence of CLL (Fig. 1F). Lastly, sorted human CD8^+^ T-cells both from healthy donors (Fig. 1G, 1I) and CLL patients (Fig. 1H,1J) also show susceptibility to IL-10. When stimulated with plate-bound anti-CD3 and soluble anti-CD28, exogenous IL-10 treatment decreased human CD8^+^ T-cell proliferation (Fig. 1G-H), IFN-γ expression (Fig. 1I-J), and CD69 expression (Fig. S1A-B) in a dose-dependent manner. 1μM MTM_ox_32*E* treatment did not dampen healthy donor nor CLL patient human CD8^+^ T-cell responses to stimuli *ex vivo* (Fig. S1C-F), suggesting this analogue could suppress CLL IL-10 without decreasing T-cell functionality.

### MTM_ox_32*E* suppresses IL-10 and preserves T-cell function

Dose response studies showed that MTM_ox_32*E* decreased Eμ-TCL1 IL-10 production with an IC50 of 70nM (Fig. 2A). In contrast, MTM_ox_32*E* treatment did not significantly inhibit cytokine production by isolated murine CD4^+^ or CD8^+^ T-cells in 24-hour cultures (Fig. 2A), or CD8^+^ T-cell proliferation in 48-hour cultures (Fig. 2B). MTM_ox_32*E* decreased hCLL PBMC IL-10 production in 93% of patients tested (Fig. 2C, Table S3) and suppressed IL-10 production by the human CLL cell line, Mec-1 (Fig. S1G). Chromatin immunoprecipitation with Sp1 in Mec-1 cells showed MTM_ox_32*E* reduced Sp1 occupancy of two GC rich sites in the IL-10 promoter (Fig. 2D) and reduced IL-10 transcription (Fig. S1H), consistent with Sp1 displacement. Notably, the mRNA levels of Sp1 target genes involved in cell cycle and survival were less affected by MTM_ox_32*E* than MTM in hCLL cells (Fig. S1I), leading to less inhibition of cellular viability (Fig. S1-J-L). Relative to the parent molecule MTM, MTM_ox_32*E* is similarly potent at suppressing Eμ-TCL1 IL-10 (Fig. S2A) but is significantly less inhibitory to murine T-cell cytokine production (Fig. S2B) and viability (Fig. S2C-F). MTM_ox_32*E* only mildly affected the viability of healthy human PBMCs after five days in culture (Fig. S3A) and did not decrease human CD4^+^ and CD8^+^ T-cell cytokine production in mixed cultures after 5 days at the dose (100nM) that blocks IL-10 production (Fig. S3B-C), though there was some inhibition at higher doses (data not shown). Taken together, these results indicate MTM_ox_32*E* reduces CLL-derived IL-10 production while preserving T-cell functionality.

**Figure 2:**
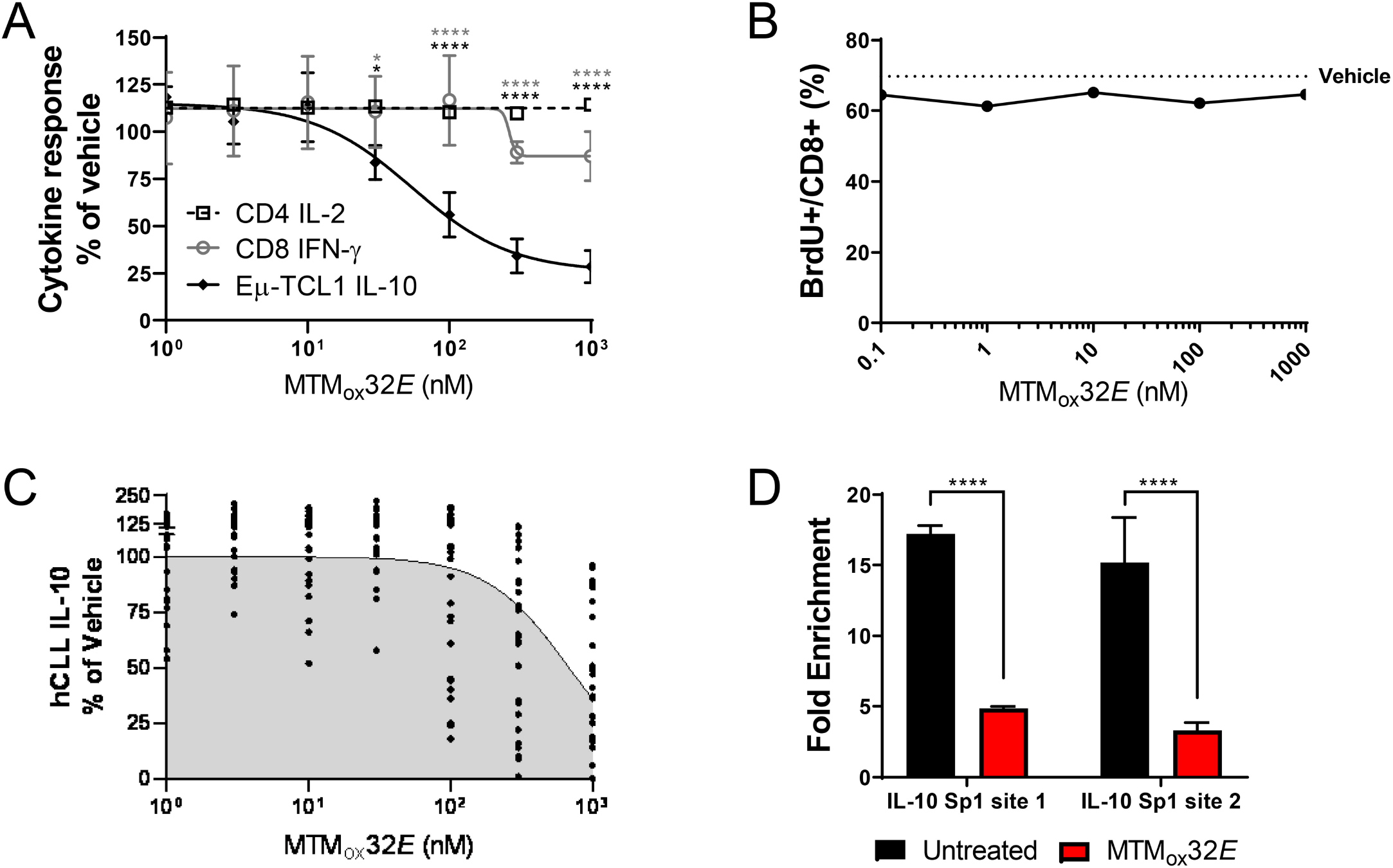
MTM_ox_32*E* suppresses CLL IL-10 without dampening *in vitro* T-cell responses. (A) Secreted IL-10 from Eμ-TCL1 splenocytes, IL-2 from murine C57BL/6 CD4^+^ T-cells, and IFN-γ from murine C57BL/6 CD8^+^ T-cells cultured for 24 hours with MTM_ox_32*E*, normalized to vehicle control (660pg/mL, 398pg/mL, 106pg/mL, respectively). T-cells were stimulated with 10μg/mL soluble anti-CD3. (B) Proliferation (BrdU incorporation) of C57BL/6 CD8^+^ T-cells cultured for 72 hours with 10μg/mL anti-CD3 ± MTM_ox_32*E* (error bars are too small to be seen). (C) Secreted IL-10 from human CLL PBMCs stimulated with 25μg/mL anti-IgM for 24 hours with MTM_ox_32*E*, normalized to vehicle control (17-656pg/mL). (D) Promoter occupancy of Sp1 GC-rich sites on the human IL-10 promoter in Mec1 cells treated with 1000nM MTM_ox_32*E*. Statistical comparisons made by two-way ANOVA in A and one-way ANOVA in D. *p<0.05, **p<0.01, ***p<0.001, ****p<0.0001

### Inhibiting CLL-derived IL-10 production enhances T-cell antitumor immunity

Since anti-IL-10 and MTM_ox_32*E* similarly suppressed CLL IL-10 and enhanced primed CD8+ T-cell activity in co-culture, these treatments were tested in the adoptive transfer model of Eμ-TCL1 in NSG mice(4). Disease burden was monitored by the appearance of CD5^+^CD19^+^ CLL cells in the blood and mice were euthanized when blood CLL reached 70-80% (Fig. S4A, S5A). Without T-cells, CLL grew to 70-80% of peripheral blood cells in 15-16 days (Fig. 3A-B). Transferring primed CD8+ T-cells at 1:32 ratio (CD8^+^:CLL) significantly delayed the development of disease, and suppressing IL-10 further delayed disease development (Fig. 3A-B). When mice were treated with anti-IL-10, leukemic burden is reduced in blood (Fig. 3A), spleen and bone marrow (Fig. S4B-C). Similarly, 12mg/kg MTM_ox_32*E* slowed disease progression in the blood, with an earlier reduction in CLL burden (statistically significant decrease at day 14 for MTM_ox_32*E* but not anti-IL-10, Fig. 3A-B) and less variability in the spleen and bone marrow (Fig. S4B-C, S5B-D). Both treatments reduced plasma IL-10 levels (Fig. 3C-D), demonstrating that MTM_ox_32*E* can suppress CLL-derived IL-10 *in vivo*, leading to decreased CLL burden.

**Figure 3:**
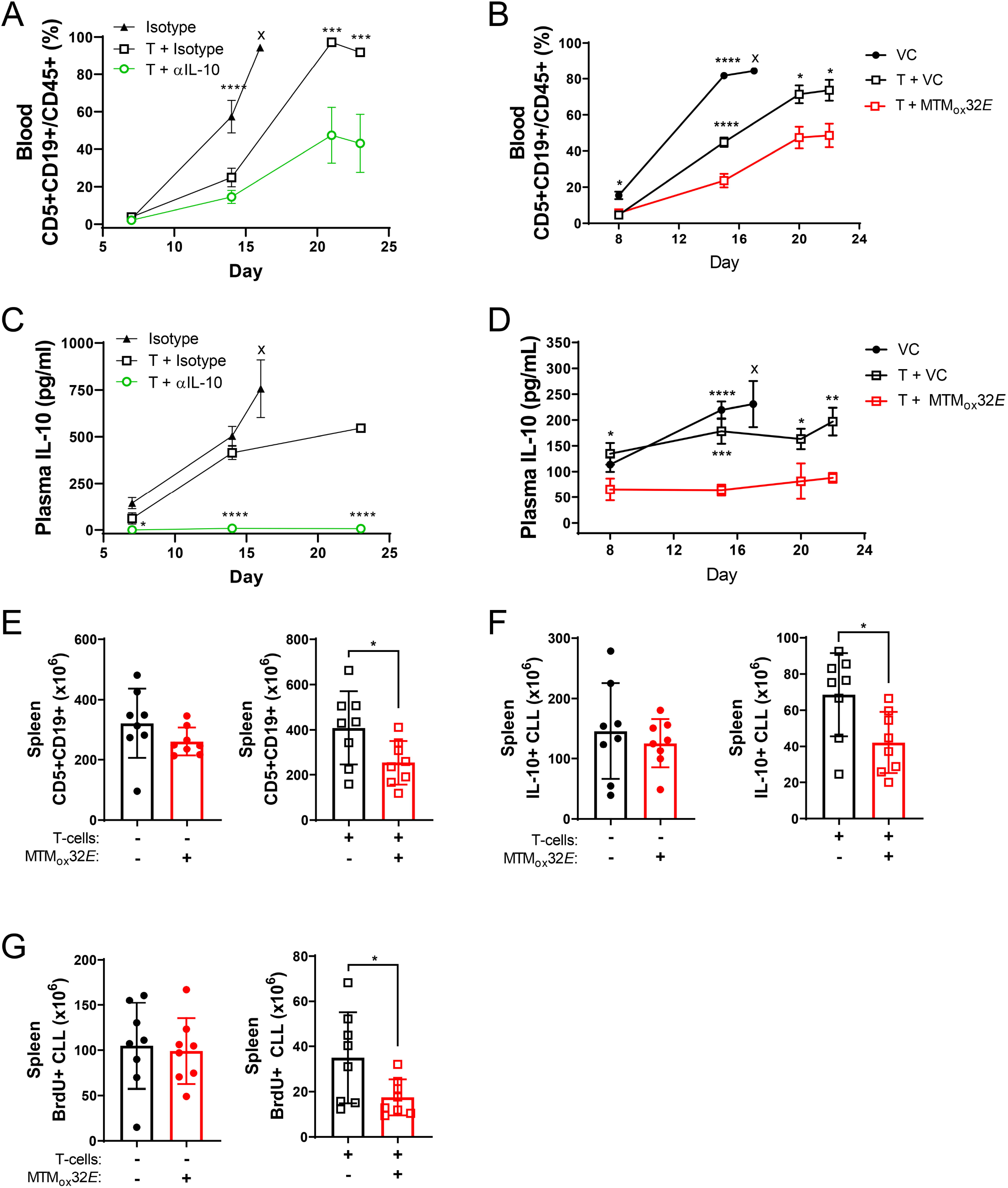
MTM_ox_32*E* enhances anti-CLL activity of CD8^+^ T-cells and suppresses CLL IL-10 production *in vivo*. (A-B) Frequency of Eμ-TCL1 CLL cells in the blood of NSG mice (mean + SEM). Six groups of eight NSG mice were injected with Eμ-TCL1 CLL, with or without Eμ-TCL1 primed CD8^+^ T-cells (CLL to T-cell ratio of 32:1) and received 150μg anti-IL-10 or isotype every two days (A) or 12mg/kg MTM_ox_32*E* or vehicle every three days (B). Stars indicate statistical significance of differences between control and anti-IL-10 (A) or MTM_ox_32*E* (B) treated group. (C-D) Plasma IL-10 levels in NSG mice (mean + SEM). Stars indicate significance between control and anti-IL-10 (C) or MTM_ox_32*E* (D) treated group. (E) Burden of Eμ-TCL1 CLL cells in the spleens of NSG mice without CD8^+^ T-cells (left panel, euthanized day 17) and with CD8^+^ T-cells (right panel, euthanized day 22) in the MTM_ox_32E group. (F) Count of IL-10^+^ Eμ-TCL1 CLL cells in the spleens of NSG mice without CD8^+^ T-cells (left, day 17) and with CD8^+^ T-cells (right, day 22). (G) Count of proliferative (BrdU^+^) Eμ-TCL1 CLL cells in the spleens of NSG mice without CD8^+^ T-cells (left, day 17) and with CD8^+^ T-cells (right, day 22). Statistical comparisons made by two-way ANOVA in A-D, and one-way ANOVA in E-G. *p<0.05, **p<0.01, ***p<0.001, ****p<0.0001, VC=vehicle control, x=mice euthanized at indicated earlier timepoint

Spleens from NSG mice also showed decreased CLL burden when MTM_ox_32*E* suppressed IL-10. Groups without T-cells had to be euthanized earlier due to faster CLL progression (day 17 vs 22), so comparisons are made within T-cell and no T-cell groups. MTM_ox_32*E* treatment reduced CLL cell frequency in the spleen and bone marrow (Fig.S5B-D). However, splenic CLL burden (count) was only reduced in NSG mice given primed CD8^+^ T-cells and MTM_ox_32*E* treatment (Fig. 3E). The number of IL-10+ splenic CLL cells (Fig. 3F, Fig. S5E) and proliferative CLL cells (Fig. 3G) were also only reduced by IL-10 suppression when T-cells were present, indicating a requirement for T-cells. The lack of significance without T-cells may be because IL-10 does not affect CLL directly but inhibits the anti-CLL T-cell response.

### *In vivo* T-cell activation increases with IL-10 suppression

When IL-10 was suppressed with MTM_ox_32*E*, CD8^+^ T-cell functionality improved in Eμ-TCL1 recipient NSG mice. CD8^+^ T-cells persisted throughout the experiment, with an increased count in the blood and a trend towards increase in the spleen after MTM_ox_32E treatment (Fig. 4A, Fig. S5F-G). Like anti-IL-10 (Fig. S4D-E), MTM_ox_32*E* treatment increased CD8^+^ T-cell proliferation, shown by increased number and frequency of splenic BrdU^+^CD5^+^CD19^-^ T-cells (Fig. 4B). On average, the number and frequency of IFN-γ^+^CD8^+^ (Fig. 4C) and Granzyme-B^+^CD8^+^ (GzB^+^CD8^+^) splenic T-cells (Fig. 4D) increased with MTM_ox_32*E* treatment. Some mice upregulated cytokine production more than others, but mice with more proliferative T-cells also had more cytokine competent T-cells. With IL-10 suppression the frequency of CD8^+^ cells expressing the effector T-cell marker, killer cell lectin-like receptor G1 (KLRG1), and the memory marker CD27 (Fig. 4E) increased. KLRG1+ effector T-cells are more cytotoxic and enhance protective immunity, and may be more active in control of CLL(29). Despite these changes, T-cells increased PD-1 expression with time, a marker of activation and eventual exhaustion, regardless of drug treatment (Fig. 4F). Therefore, we hypothesized anti-CLL immunity could be improved by combining IL-10 suppression with ICB.

**Figure 4:**
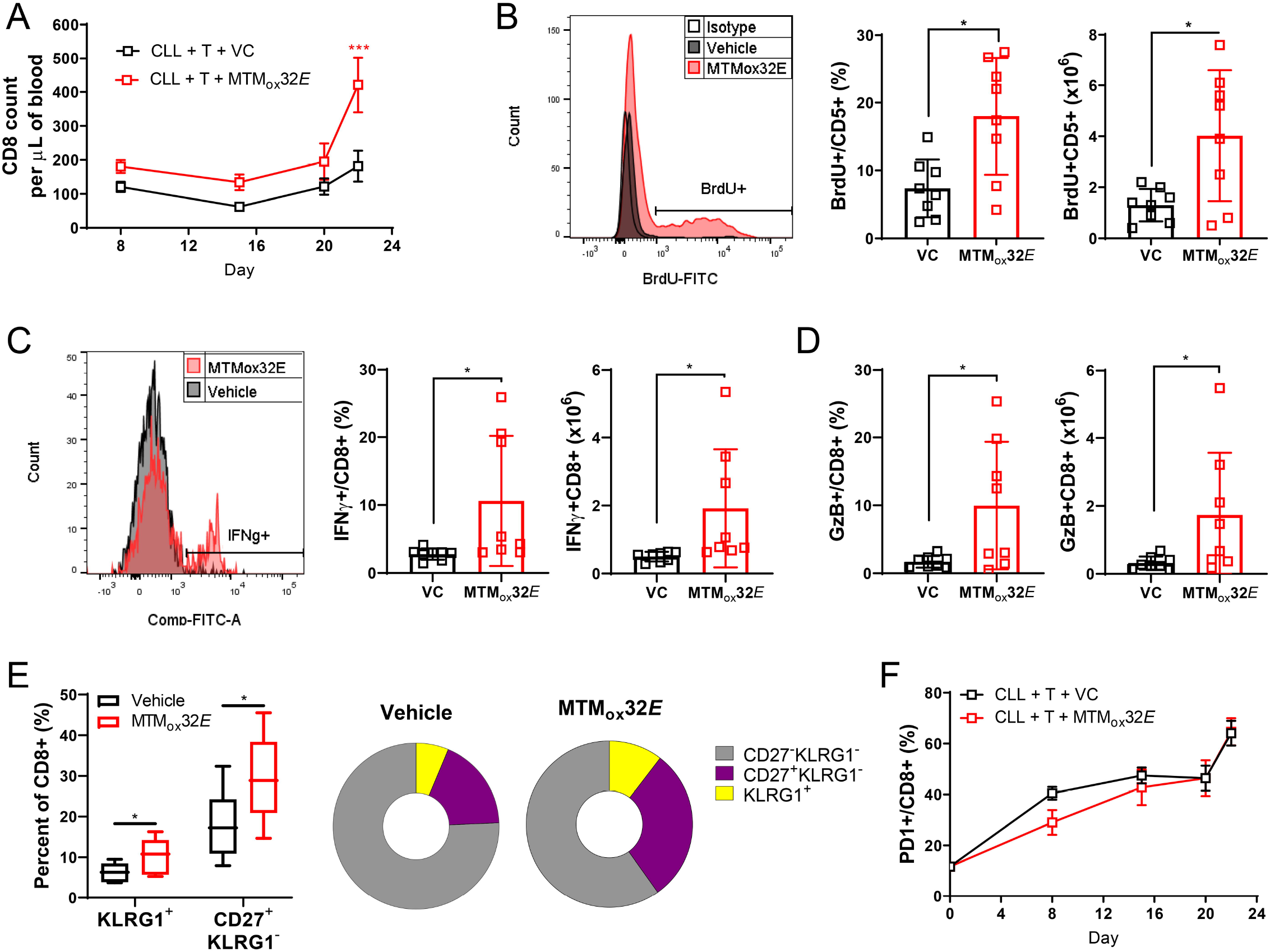
Inhibiting CLL IL-10 with MTM_ox_32*E* increases the numbers of T-effector cells. Mice were treated as in Figure 3B. (A) The number of CD8^+^ T-cells in the blood of NSG mice. (B) (Left) Representative histograms of BrdU incorporation by splenicCD5^+^CD19^-^ T-cells (>99% CD8^+^). Frequency (middle) and number (right) of BrdU^+^ splenic CD8^+^ T-cells. (C) (Left) Representative histograms of IFN-γ production by splenic CD8^+^ T-cells. Frequency (middle) and number (right) of splenic IFN-γ^+^ CD8^+^ T-cells. (D) Frequency (left) and number (right) of GzB^+^ CD8^+^ T-cells in the spleen. (E) Frequency of KLRG1^+^ highly cytotoxic effector and CD27^+^ memory CD8^+^ T-cells in the spleen (left) and pie chart representations of CD8^+^ T-cell proportions. (F) Frequency of PD-1 expressing CD8^+^ T-cells in the blood. Statistical comparisons in A+F were obtained by two-way ANOVA comparing vehicle to MTM_ox_32*E* treated mice, and in B-E by one-way ANOVA between vehicle to MTM_ox_32*E*. *p<0.05

### Inhibiting CLL-derived IL-10 production improves responses to anti-PD-L1 checkpoint blockade

Previous reports described ICB reducing CLL burden in Eμ-TCL1 mice, especially in combination with other therapies(13, 30). Our results were similar, where anti-PD-L1 ICB alone modestly reduced CLL prevalence in blood and bone marrow but not spleen (Fig. S6A-D). Although there were more proliferating CD8^+^ T-cells in spleen (Fig S6E), ICB did not alter T-cell abundance, and yielded fewer IFN-γ^+^ and more PD-1^+^CD8^+^ T-cells (Fig. S6F-G). We tested if IL-10 suppression could enhance the efficacy of anti-PD-L1 ICB in NSG mice given purified CD19^+^ Eμ-TCL1 CLL cells and primed CD8^+^ T-cells.

With anti-PD-L1 ICB and MTM_ox_32*E*-mediated IL-10 suppression combined (Fig.S7A), antitumor CD8^+^ T-cells were more effective at controlling Eμ-TCL1 CLL growth. Combination therapy slowed CLL growth in the blood compared to anti-PD-L1 alone or isotype plus vehicle treated control mice (Fig. 5A). By the end of the experiment, CLL frequency (Fig. 5B) and burden (Fig. 5C) in the spleen were on average 4.5-fold lower in combination therapy mice versus anti-PD-L1 alone (68.9% vs 15.4%; 94.9×10^6^ vs 21.1×10^6^ cells). In the spleen, the frequency and number of CD8^+^ T-cells more than doubled (combination vs ICB: 6.97% vs 15.4%, 9.81×10^6^ vs 22.14×10^6^ cells, Fig. 5D). Blood CD8^+^ T-cell levels were also elevated with combination therapy (Fig. 5E, Fig. S7B). Spleen size was not different between the therapy groups, which could be due to the expansion of T-cells with combination therapy (Fig. S7C-D). Combination therapy reduced CLL spread to the bone marrow (combination vs ICB: 1.58% vs 0.28%, 1.6×10^5^ vs 0.2×10^5^, Fig. 5F), and dramatically altered the severity of disease, as only one mouse receiving combination therapy developed more than 30% CLL in the blood by the end of the experiment (Fig. 5G).

**Figure 5:**
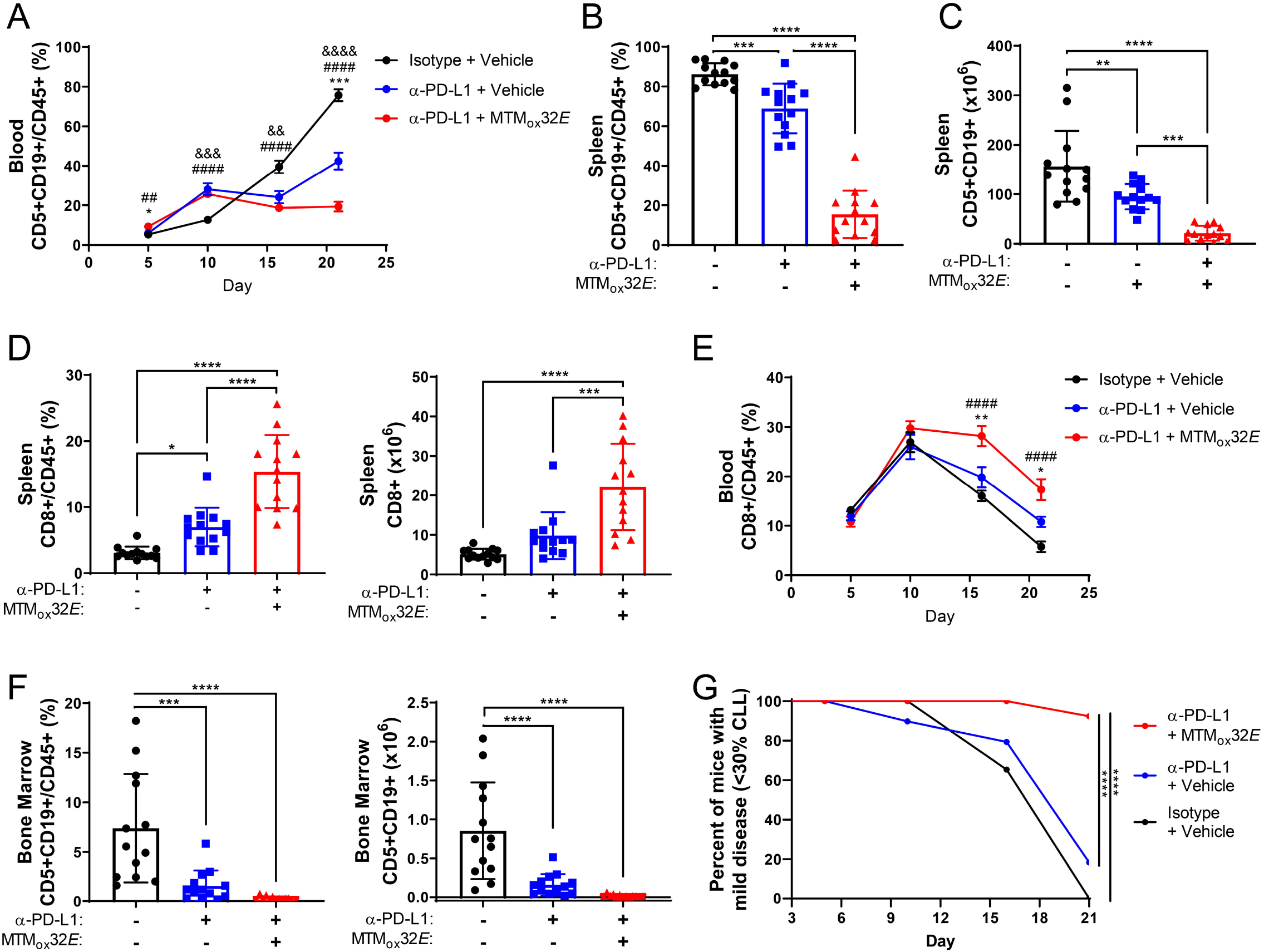
Anti-PD-L1 checkpoint blockade is more effective when combined with IL-10 suppression by MTM_ox_32*E*. Three groups of thirteen NSG mice were injected with Eμ-TCL1 and Eμ-TCL1 primed CD8^+^ T-cells at a ratio of one T-cell to 32 CLL cells. Mice received 12mg/kg MTM_ox_32*E* or vehicle every two to three days plus 10mg/kg anti-PD-L1 or isotype control every three days. (A) Frequency of Eμ-TCL1 CLL cells in the blood of NSG mice (mean + SEM). (B-C) Frequency (B) and count (C) of Eμ-TCL1 CLL cells in the spleen. (D) Frequency (left) and count (right) of CD8^+^ T-cells in the spleen of recipient NSG mice. (E) Frequency of CD8^+^ T-cells in the blood of NSG recipients (mean + SEM). (F) Frequency (left) and count (right) of Eμ-TCL1 CLL cells in the bone marrow. (G) Percent of mice with less than 30% CLL in the blood over time. Statistical significance in A, E, G was calculated by two-way ANOVA comparing vehicle to treated mice, and by one-way ANOVA comparing vehicle to treated mice in remaining panels. *p<0.05, **p<0.01, ***p<0.001, ****p<0.0001 indicate statistical significance of *difference between combination and ICB alone, #difference between combination and control, &difference between ICB alone and control

Adding IL-10 suppression to ICB altered the proliferative capacity of both CD8+ T-cells and CLL cells in NSG mice. As expected, PD-L1^+^ CLL cells were decreased with ICB treatment (Fig. S7E). Plasma IL-10 levels were lowest in mice receiving combination therapy, with intermediate IL-10 levels for anti-PD-L1 alone (Fig. S7F). The number of BrdU^+^CD8^+^ T-cells were elevated in both groups (Fig. 6A), but the number of dividing CLL cells only decreased with combination therapy (Fig. 6B). This resulted in a greatly elevated ratio of total and proliferative CD8^+^ to CLL cells with combination therapy (Fig. S7G). Interestingly, plasma IL-10 levels inversely correlated with the number of total and dividing CD8^+^ T-cells, suggesting IL-10 plays a role in decreasing T-cell persistence and proliferation (Fig. S7H).

**Figure 6:**
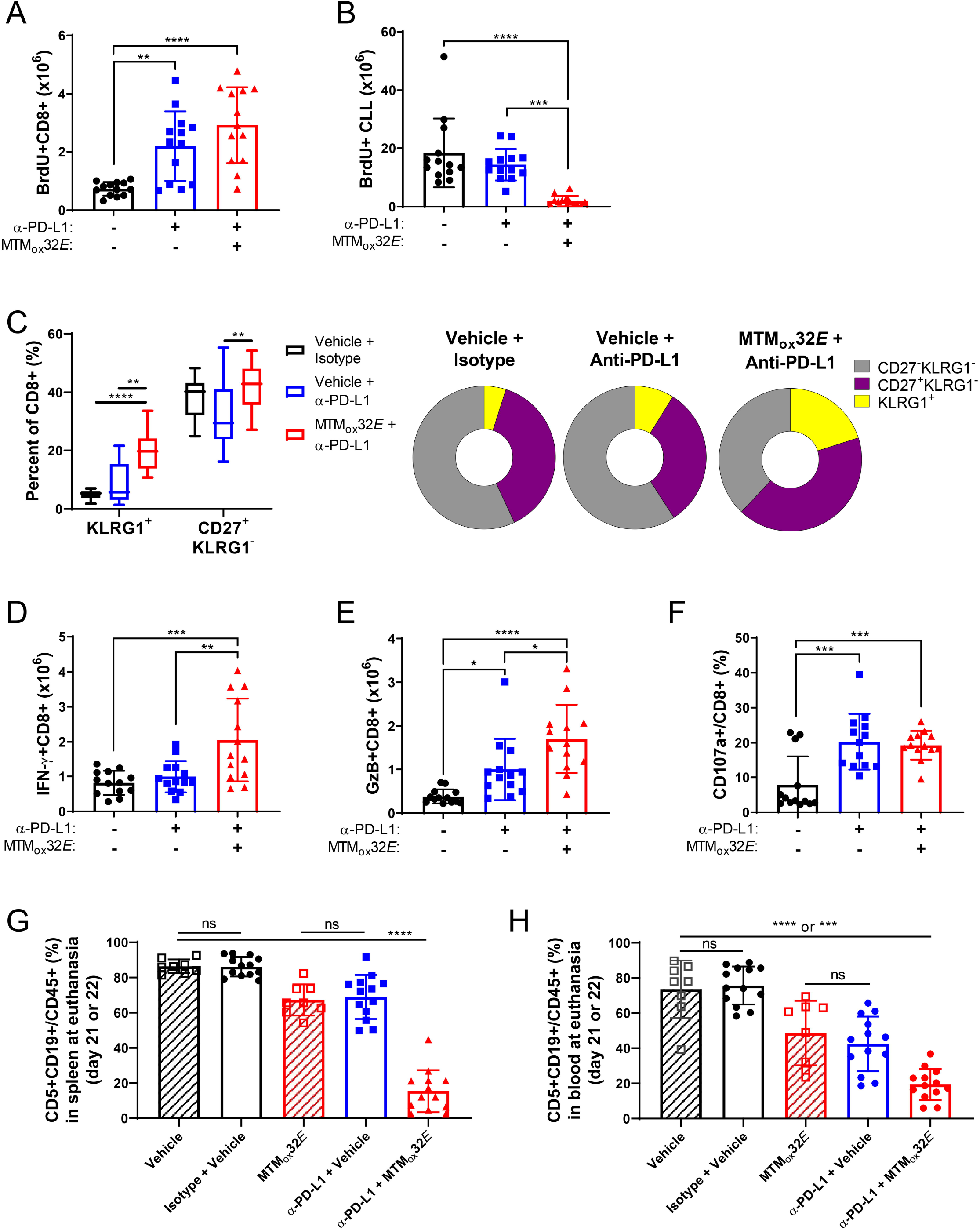
T-cells from Eμ-TCL1 mice are more functional when treated with both anti-PD-L1 checkpoint blockade and IL-10 blockade. Mice were treated as in Figure5. (A) Count of BrdU^+^ splenic CD8^+^ T-cells. (B) Count of BrdU^+^ Eμ-TCL1 CLL cells in the spleen. (C) Frequency of KLRG1^+^ highly cytotoxic effector and CD27^+^ memory CD8^+^ T-cells in the spleen (left) and pie chart representations of CD8^+^ T-cell proportions. (D-E) Count of IFN-γ^+^ (D) and GzB^+^ (E) splenic CD8^+^ T-cells. (F) Frequency of CD107a^+^ splenic CD8^+^ T-cells. (G) Comparison of CLL frequency in spleen from Fig. 5B and Fig. S5B. (H) Comparison of CLL frequency in the blood from Figures 3B and 5A. Statistical comparisons done by one-way ANOVA comparing vehicle to treated mice, except in G+H where all groups were compared to each other. *p<0.05, **p<0.01, ***p<0.001, ****p<0.0001

When combining IL-10 suppression and ICB therapy, the outcome of T-cell activation was further altered. MTM_ox_32*E* treatment increased the proportion of highly cytotoxic effector T-cells (KLRG1^+^) and memory T-cells (CD27^+^) over ICB alone (Fig. 6C). There were also higher numbers of splenic IFN-γ^+^CD8^+^ and GzB^+^CD8^+^ T-cells (Fig. 6D-E), and CD8^+^ T-cells expressing CD107a, a marker of degranulation, in treated mice (Fig. 6F). Still, similar levels of PD-1^+^CD8^+^ T-cells were detected in the blood throughout the experiment (Fig. S8A). PD-1 is upregulated after T-cell activation, and truly exhausted T-cells often express other exhaustion markers in addition to PD-1 (Fig. S8B). Though there was a higher frequency of splenic PD-1^+^CD8^+^ T-cells in combination therapy mice (Fig. S8C), very few CD8^+^ T-cells expressed other markers of exhaustion, such as CTLA4, Lag3 or Tim-3 (Fig. S8D). The total frequency of exhausted cells (PD-1^+^CTLA4^+^, PD-1^+^Lag3^+^, or PD-1^+^Tim-3^+^) decreased for both treatment groups (Fig. S8E), suggesting that most of the PD-1+ T-cells were not yet exhausted.

The increase in T-cell functionality with IL-10 suppression and ICB combination therapy shows clearly enhanced antitumor immunity, so lastly, we compared CLL burden within the previous experiment. With this comparison, the additive effect of IL-10 suppression and ICB became clear. CLL frequency is markedly lower in the spleen (Fig. 6G) and blood (Fig. 6H) with the combination therapy compared to either control or monotherapy. Furthermore, when mice are treated with this combination, median survival increases significantly from 41 to 49.5 days (Fig. S8F). Taken together, these data show improved CD8^+^ T-cell antitumor immunity in CLL with combined ICB and IL-10 suppression.

## Discussion

IL-10 is a potent anti-inflammatory cytokine that downregulates CD8^+^ antitumor immunity(4). In our studies the therapeutic suppression of IL-10 enhanced CD8^+^ T-cell control of CLL. IL-10 suppression was most effective at enhancing anti-CLL CD8^+^ T-cell activity when combined with anti-PD-L1 ICB. This combination reduced disease burden compared to ICB alone and increased the number of CD8^+^ T-cells, suggesting improved persistence. The outcome of T-cell activation changed as the numbers of CD8^+^ T-cells expressing functional markers like IFN-γ, GzB, CD27, and KLRG1 increased and exhausted CD8^+^ T-cells decreased. These data show IL-10 suppression can improve CD8^+^ T-cell functionality, antitumor immunity, and responses to ICB. They point to IL-10 suppression as a novel therapeutic strategy that may improve responses to T-cell-based immunotherapy in human CLL.

In CLL and several solid tumors, IL-10 reduced the inflammatory state of immune responses including CD8^+^ T-cells(23, 31-33), although there are examples where IL-10 supported antitumor T-cell function(34-36). The sources of IL-10 are diverse ranging from T-reg cells, macrophages, plasma cells, B-reg cells and cancer cells, as in the present case. Indeed, the role of IL-10 in cancer may be context dependent. In liver and prostate cancers, IL-10 inhibits cytotoxic T-cell function and anti-cancer immunity(37, 38). PEGylated IL-10 treatment increased STAT1 and STAT3 activation in CD8^+^ T-cells, enhancing control of PDV6 squamous carcinoma tumors(35). The dose and duration of IL-10 exposure is critical to its regulatory effects on CD8^+^ T-cell maturation into functional memory cells(39). Our data suggest that IL-10 suppression in aggressive CLL enhances memory and effector cell development from CLL-primed CD8^+^ T-cells, which may differ from T-cells that are mostly tumor-antigen naïve or exhausted. The IL-10 pathway also plays a role in T-cell exhaustion during chronic viral infection, and can directly downregulate CD8^+^ T-cell function(40, 41). Blocking IL-10 and PD-1 during chronic viral infection synergistically enhanced viral control and reversed CD8^+^ T-cell exhaustion(42). Furthermore, we previously showed IL-10 inhibits macrophage pro-inflammatory cytokine secretion in the aged, which may be important as CLL mainly affects older individuals(43). CD138^+^PD-L1^+^ plasma cell derived IL-10 also affects immune responses to infectious agents in part by inhibiting innate immune cells(44). These innate cells and stromal cells are part of the microenvironment which supports CLL cell growth. We find that coculture of CLL cells with stromal cells dramatically enhances IL-10 production by CLL cells (Collard *et al*, manuscript in preparation).

We describe MTM_ox_32*E*, a novel small molecule inhibitor of CLL cell IL-10 production, which does not inhibit effector T-cell function, and enhances anti-CLL CD8^+^ T-cell activity. MTM_ox_32*E* was more effective than anti-IL-10 in inducing a more consistent reduction in CLL growth. A small molecule may have better access than antibodies to microenvironments where CLL cells interact with other cell types. MTM_ox_32*E* treatment reduced Sp1 occupancy of the IL-10 promoter and CLL IL-10 secretion but is uniquely less toxic than MTM. Like MTM, MTM_ox_32*E* interacts with GC rich regions of DNA. MTM_ox_32*E* had better pharmacokinetics, tolerability, and selectivity than MTM and other analogues(26). MTM_ox_32*E* showed some selectivity for inhibiting Sp1 binding on the IL-10 promoter in cell culture. This may be due to interactions between the analogue and additional proteins in the transcription complex promoting CLL IL-10 production, which requires further investigation. MTM_ox_32*E* also has a small inhibitory effect on CLL growth *in vivo* but less so when used as single agent *in vitro*. This could be due to its effect on the microenvironment, which requires further investigation. Sp1 positively regulates murine T-BET (T-box expressed in T-cells), which controls natural killer and T-cell IFN-γ expression, and MTM treatment reduced both T-BET and IFN-γ production in natural killer cells(45). Sp1 expression and transcriptional activity was shown to be upregulated in activated and proliferating CD4^+^ T-cells(46, 47), but not in activated CD8^+^ T-cell subsets(48, 49). Our studies focused on CD8^+^ T-cells, where we see no detrimental effect of MTM_ox_32*E* treatment. Additional studies are required to elucidate the differential effects of MTM and MTM_ox_32*E* on naïve, effector and memory T-cells.

Expression of IL-10 by CLL cells is upregulated by B-cell receptor (BCR) signaling and may be reduced by inhibitors of this pathway. Many therapeutics for CLL inhibit kinases in BCR signaling(50, 51), and have yielded great clinical outcomes for CLL patients. The ability of Bruton’s tyrosine kinase (Btk) inhibitors (i.e. ibrutinib) and phosphoinoside-3 kinase (PI3K) inhibitors (i.e. idelalisib) to inhibit CLL growth are well studied, and may also enhance immune responses(52). However, some patients can develop resistance mutations or have worse disease after treatment cessation(53, 54). Ibrutinib treatment also affects T-cells, enhancing their function in CLL patients, which may be explained by reduced CLL-derived IL-10 in addition to its direct effects(55). CD8^+^ T-cells in hCLL exist in a pseudo-exhausted state(8), and T-cell exhaustion may occur within certain T-cell populations in patients(56). Reversing this exhaustion may be possible with the right therapeutic strategy(57). In mouse studies blocking the PD-1/PD-L1 pathway was promising, but was more effective with additional blockade of Lag-3 or Tim-3(30, 58). Anti-PD-1 ICB primarily affects CD8+ T-cells and can rescue exhausted T-cells, in contrast with anti-CTLA4(59). Nivolumab (anti-PD-1) is currently in clinical trials for human CLL in combination with the Btk inhibitor ibrutinib (NCT02420912) since previous reports suggested this combination would be more effective than monotherapy(12-14). Our findings suggest combining PD-1/PD-L1 blockade with IL-10 suppression could also improve antitumor T-cell responses (Fig. 7) and may still be effective for patients who have developed resistance to Btk inhibitors.

**Figure 7:**
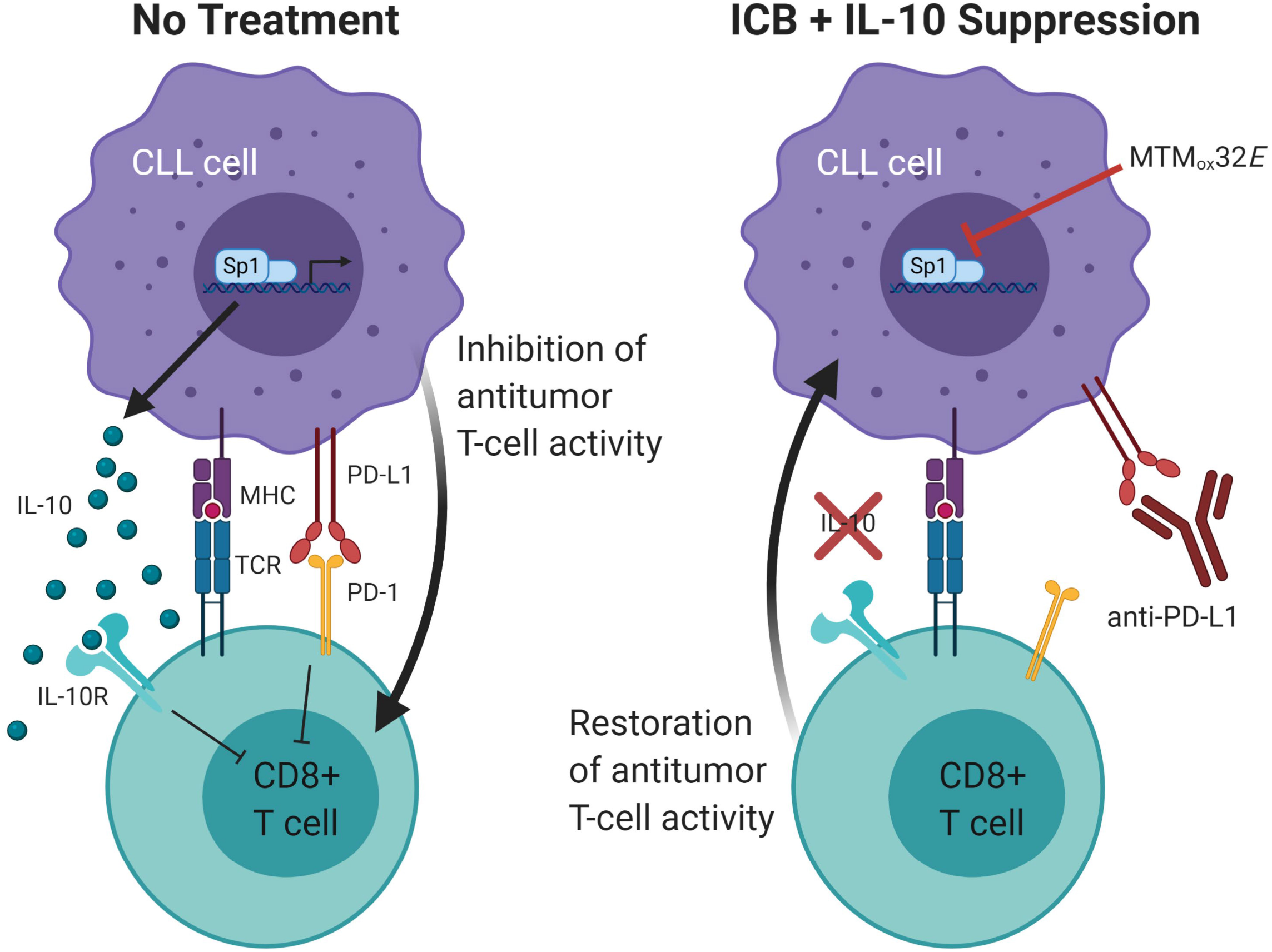
Cooperation of IL-10 suppression and immune checkpoint blockade. Created with BioRender.com. Model shows combining IL-10 suppression with anti-PD-L1 ICB enhances antitumor CD8^+^ T-cell activity by counteracting multiple methods of CLL immune suppression.

With no known cure and increasing CLL incidence(60), there is a need for novel therapies for CLL. Decreasing IL-10 to improve antitumor immunity provides an opportunity to overcome CLL-induced immunosuppression and enhance current CLL immunotherapies. Immune evasion through T-cell suppression is common to many types of cancer, so this strategy may also be applicable to other tumors(16, 19, 20, 22, 32). Further studies will reveal whether suppressing IL-10 will be effective in treating human cancer patients.

## Acknowledgements

We would like to thank the patients and their families for participating in our research and Dr. Eric Durbin with the Kentucky Cancer Registry. We also thank Dr. Siva Gandhapudi and Dylan Rivas for their helpful discussion. Jacqueline R. Rivas is a Fellow of the Leukemia & Lymphoma Society and was partially supported by the NCI Training Grant T32CA165990.

This project was supported in part by R01CA165469 (SB), R01CA217934 (SB), R01CA217255 (JST), and R01GM115261 (JST). We also acknowledge the support from the National Center for Advancing Translational Sciences (UL1TR001998 and UL1TR000117), the Center of Biomedical Research Excellence (COBRE) in Pharmaceutical Research and Innovation (CPRI, P20GM130456), the University of Kentucky College of Pharmacy PharmNMR Center, the University of Kentucky Markey Cancer Center and Markey Cancer Center’s Flow Cytometry and Immune Monitoring Shared Resource Facility (P30CA177558). The content is solely the responsibility of the authors and does not necessarily represent the official views of the NIH.

