## Supplementary Materials and Methods, Supplementary Tables 1-3, Supplementary Figures 1-8, Supplementary Figure Legends, Supplementary Table Legends for "Interleukin-10 suppression enhances T-cell antitumor immunity and responses to checkpoint blockade in chronic lymphocytic leukemia"

**Chemistry for $MTM_{ox}32E$ production**

The synthesis of $MTM_{ox}32E$ was previously described (Liu *et al*, 2020). The analogue is synthesized from MTM and characterized by mass spectrometry and determined to be greater than 95% pure by reverse phase HPLC, as detailed elsewhere. $MTM_{ox}32E$ was selected based on its improved pharmacokinetics in comparison to MTM. In comparison to other analogues, it had better selectivity efficacy and target engagement properties (Liu *et al*, 2020). The structure of $MTM_{ox}32E$ is shown below.

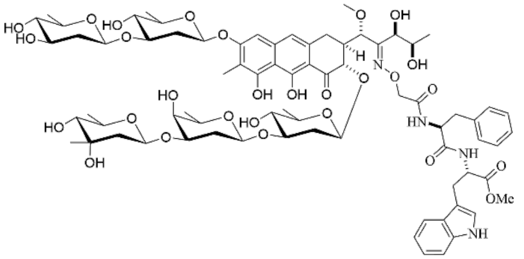

**Mouse tissue processing**

Blood obtained from live mice was done by sub-mandibular or sub-mental bleeding. After euthanizing mice, blood was collected by cardiac puncture, spleens were crushed, and bone marrow was harvested from one femur for analysis. To process the blood, 200µL was added to 5mL red blood cell lysis buffer (154.4mM $NH_4Cl$, 14.2mM $NaHCO_3$, and 1.10mM EDTA, pH 7.3), washed with HBSS and used in flow cytometry panels. The remaining blood was centrifuged for 10 minutes at max speed in a table-top Eppendorf 5430 microcentrifuge (Hamburg, Germany) to separate plasma from blood cells, and plasma was stored at -80°C. Plasma was diluted 1:3 to 1:10 with 10% FBS in PBS and IL-10 was measured with an ELISA MAX Standard Set Mouse IL-10 kit according to the manufacturer's protocol (BioLegend, San Diego, CA). Remaining spleen cells from the experiment were stored at -80°C in 10% DMSO, 90% FBS. As shown in our previous publication using this adoptive transfer system, we did not observe graft versus host disease (Alhakeem *et al*, 2018).

### Flow cytometry

Surface staining of mouse or human cells was done by blocking with 5μg normal rat IgG for mouse cells (Millipore Sigma, St. Louis, MO), or 5μg normal mouse IgG for human cells (Millipore Sigma, St. Louis, MO) for 10 minutes on ice followed by a 30-minute incubation with 1μg of each fluorophore conjugated antibody. Antibodies used in the various panels are listed in Table S1. For detection of intracellular proteins, cells were fixed with Fixation Buffer (BioLegend, San Diego, CA) and permeabilized with Intracellular Staining Perm Wash Buffer (BioLegend, San Diego, CA) according to the manufacturer's protocol, followed by a 1-hour incubation on ice with 1μg of each antibody. To detect BrdU incorporation, cells were treated with 20μg DNAse (Millipore Sigma, St. Louis, MO) for 1 hour at 37°C after permeabilizing and before antibody staining. Samples were run on an LSRII flow cytometer (BD Biosciences, Franklin Lakes, NJ) and data were analyzed with FlowJo v10 software (Tree Star, Inc., Ashland, OR).

### Healthy Human PBMC or T-cell cultures

Peripheral blood mononuclear cells (PBMCs) were isolated from healthy donors' blood (n=4, Fig. S3) or leukopaks (n=4, Fig. 1, Fig. S1) after obtaining with IRB approval. PBMCs were cultured at a concentration of $10^6$/mL in Immunocult XF-T-cell expansion medium (StemCell Technologies, Vancouver, Canada), supplemented with 5% FBS, and MACS GMP ExpAct Treg immunobeads (Miltenyi, 2 beads/cell) in the absence (control) or presence of $MTM_{ox}32E$ at the indicated doses. CD8+ T-cells were isolated from Leukopaks after Ficol-Paque density centrifugation with Miltenyi MACS CD8 MicroBeads, human and separated on a Miltenyi AutoMACS Pro. Apoptosis assays were performed after 72-hour culture with the APC Annexin V apoptosis detection kit with Propidium Iodide (PI) (BioLegend, USA) following the manufacturer instructions. For detection of intracellular levels of IFN-γ in Fig. S3, 24-hour cultured cells were restimulated for additional 4 hours with Cell Stimulation Cocktail and Protein Inhibitor Cocktail (both from eBioscience, San Diego, CA). Cells were then harvested, and single cell suspensions stained with FITC-anti-CD4 (eBioscience) and APC-Fire750 anti-CD8 (BioLegend) antibodies. Cells were fixed and permeabilized with the FoxP3 Fix/perm Buffer Set (BioLegend) and stained with eFluor450-labeled anti-IFNγ (eBioscience). Analyses were performed in a LSRII cytometer (BD Biosciences) and FlowJo software (Tree Star, Inc.). For CD8+ T-cell cultures in Fig. 1 and Fig. S1, 10μg/mL low-endotoxin azide free (LEAF) anti-human CD3 (BioLegend) was coated on the bottom of 96 well plates for 24-48 hours at 37°C. Plates were aspirated and purified CD8+ T-cells added at $1x10^6$/mL with 1μg/mL soluble LEAF anti-human CD28 (BioLegend) and varying doses of recombinant human IL-10 (Peprotech or BioLegend) and cultured for five days. In the last 6 hours, 40uM BrdU was pulsed into cultures before cells were stained with BioLegend anti-CD45 APC/Cy7, anti-CD8 APC, anti-CD69 PE/Cy7, fixed with 4% paraformaldehyde and permeabilized with BioLegend Perm Wash Buffer in their 96 well plates. Cells were then DNAse digested for 1 hour at 37°C (according to BioLegend protocol) and stained with anti-IFNγ Pacific Blue and anti-BrdU FITC. Data were acquired on a BD BioSciences Symphony A3 using the HTS plate adapter.

**Mec-1 Cell Culture**

Mec-1 cells (ATCC 74155, Manassas, VA) were cultured in complete IMDM in T75 flasks (Corning, Corning, NY) before use in assays. Cells were validated to be mycoplasma free in 2016. For secreted IL-10, $5x10^4$ cells were placed in flat bottom 96 well plates with drug concentrations indicated in Fig. S3 for 24 hours at 37°C. The amount of IL-10 was measured with an ELISA MAX Standard Set Human IL-10 kit (BioLegend, San Diego, CA).

**Chromatin Immunoprecipitation**

$10x10^6$ Mec-1 cells were treated with 1000nM $MTM_{ox}32E$ or vehicle control for 24 hours before harvest. The Simple ChIP Kit with Magnetic Beads was used according to the manufacturer's protocol (Cell

Signaling Technologies, Danvers, MA), with an anti-Sp1 antibody (product number sc-17824, Santa Cruz Biotechnology Inc., Dallas, TX) and custom IL-10 promoter primers (Integrated DNA Technologies, Coralville, IA). Primer set 1 (-1158 to -1009): FWD: AACTGGCTCCCCTTACCTTC, REV: GGCTGGATAGGAGGTCCCTT; Primer set 2 (-121 to -7): FWD: AGAAGGAGGAGCTCTAAGCA, REV: TCACCTCTCTGTCCCCCTTTTA.

**Real-time PCR**

$10x10^6$ primary hCLL PBMCs were cultured with 100nM MTM or $MTM_{ox}32E$ for 24 hours before harvest, and $5x10^6$ Mec-1 cells were cultured with 1000nM MTM or $MTM_{ox}32E$ for 24 hours before harvest. RNA was isolated with TRI Reagent according to the manufacturer's protocol (Millipore Sigma, St. Louis, MO) and stored at -80°C until use. Complimentary DNA was synthesized with the iScript cDNA Synthesis Kit (Bio-Rad Laboratories, Hercules, CA) and mRNA was quantified with iTaq Universal SYBR Green Supermix (Bio-Rad Laboratories, Hercules, CA). Samples were run on a CFX96 Real-Time PCR Detection System (Bio-Rad Laboratories) using custom primers (available upon request, Integrated DNA Technologies). Data were analyzed using Bio-Rad CFX Manager software (Bio-Rad Laboratories).

### Supplementary Figure 1

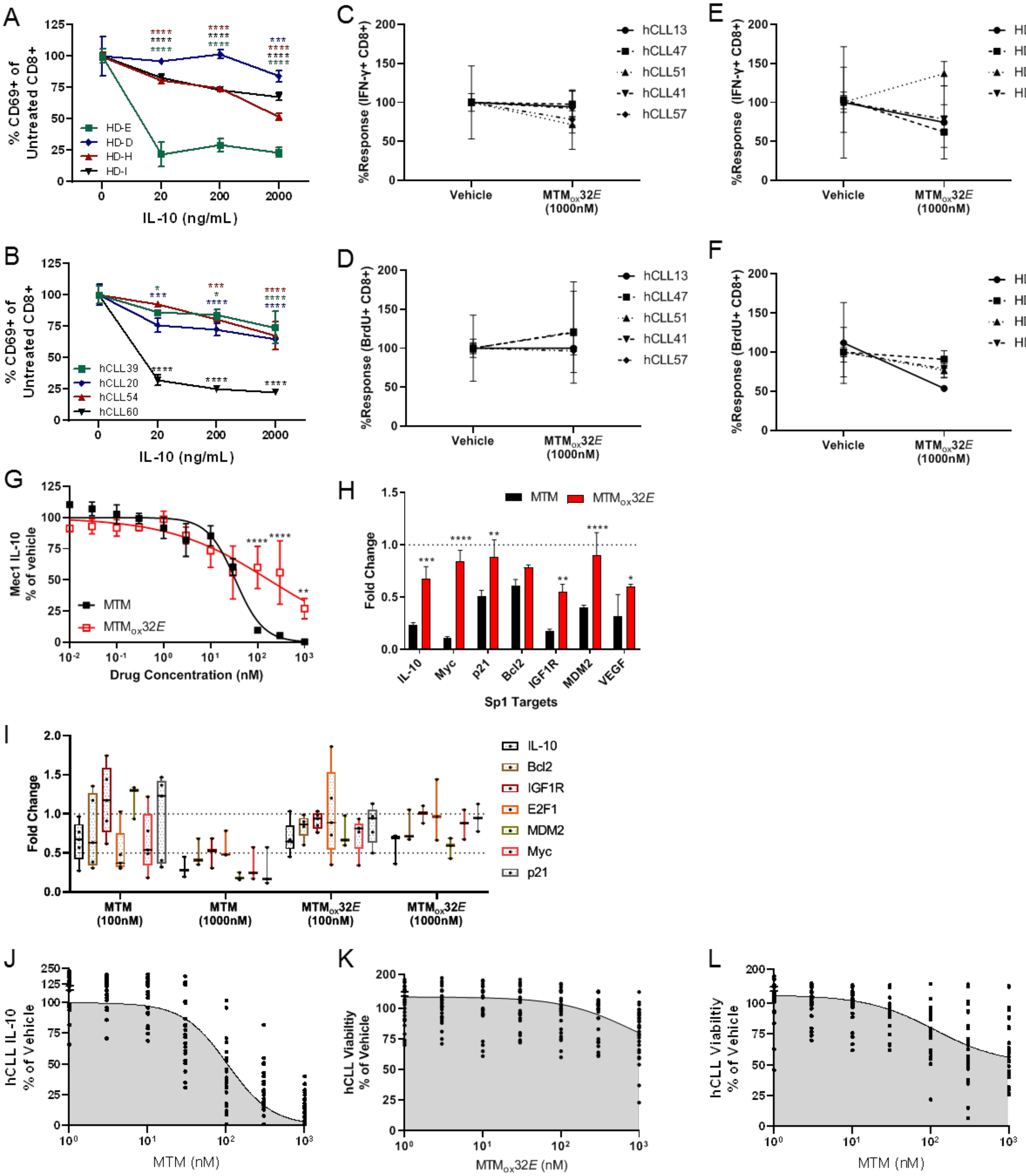

**Supplementary Figure 2**

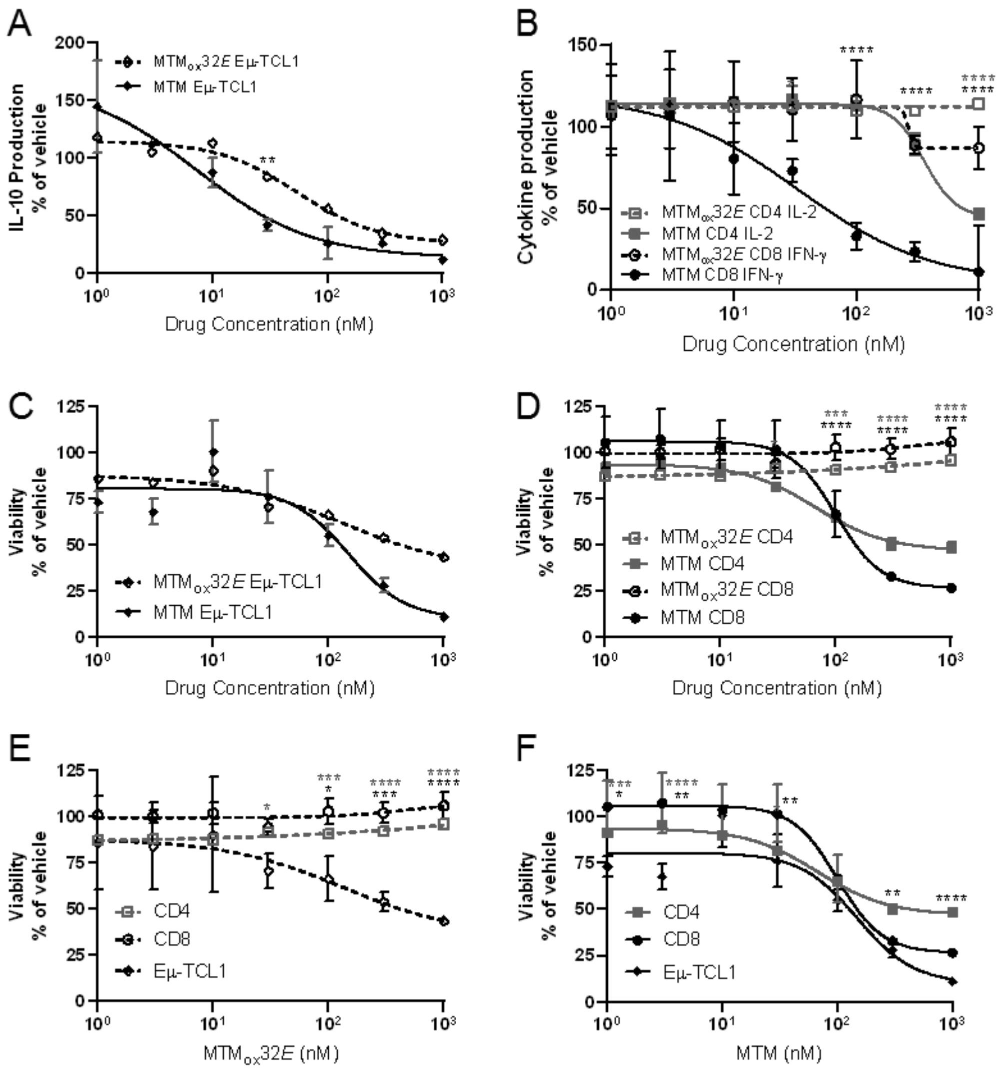

**Supplementary Figure 3**

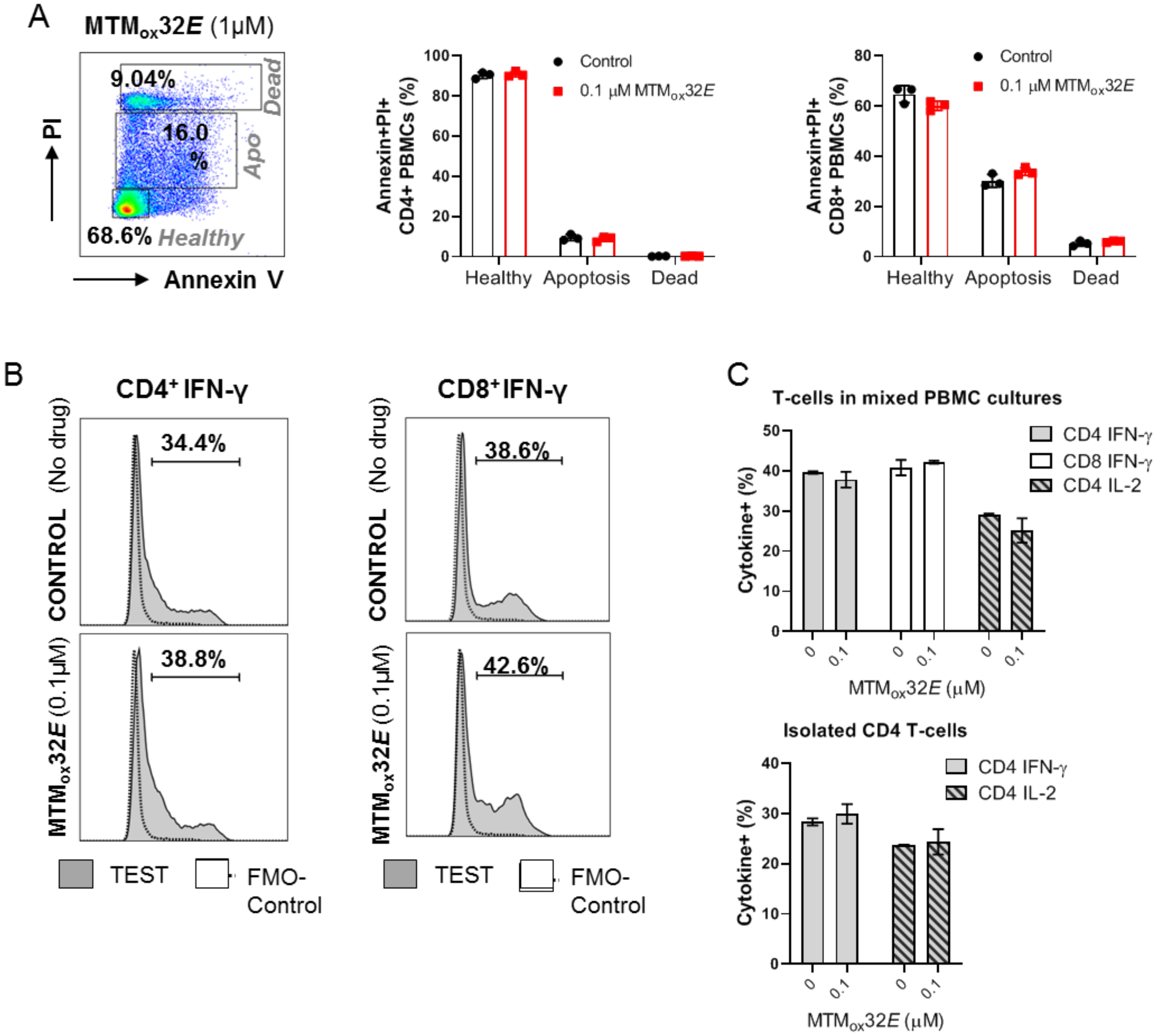

**Supplementary Figure 4**

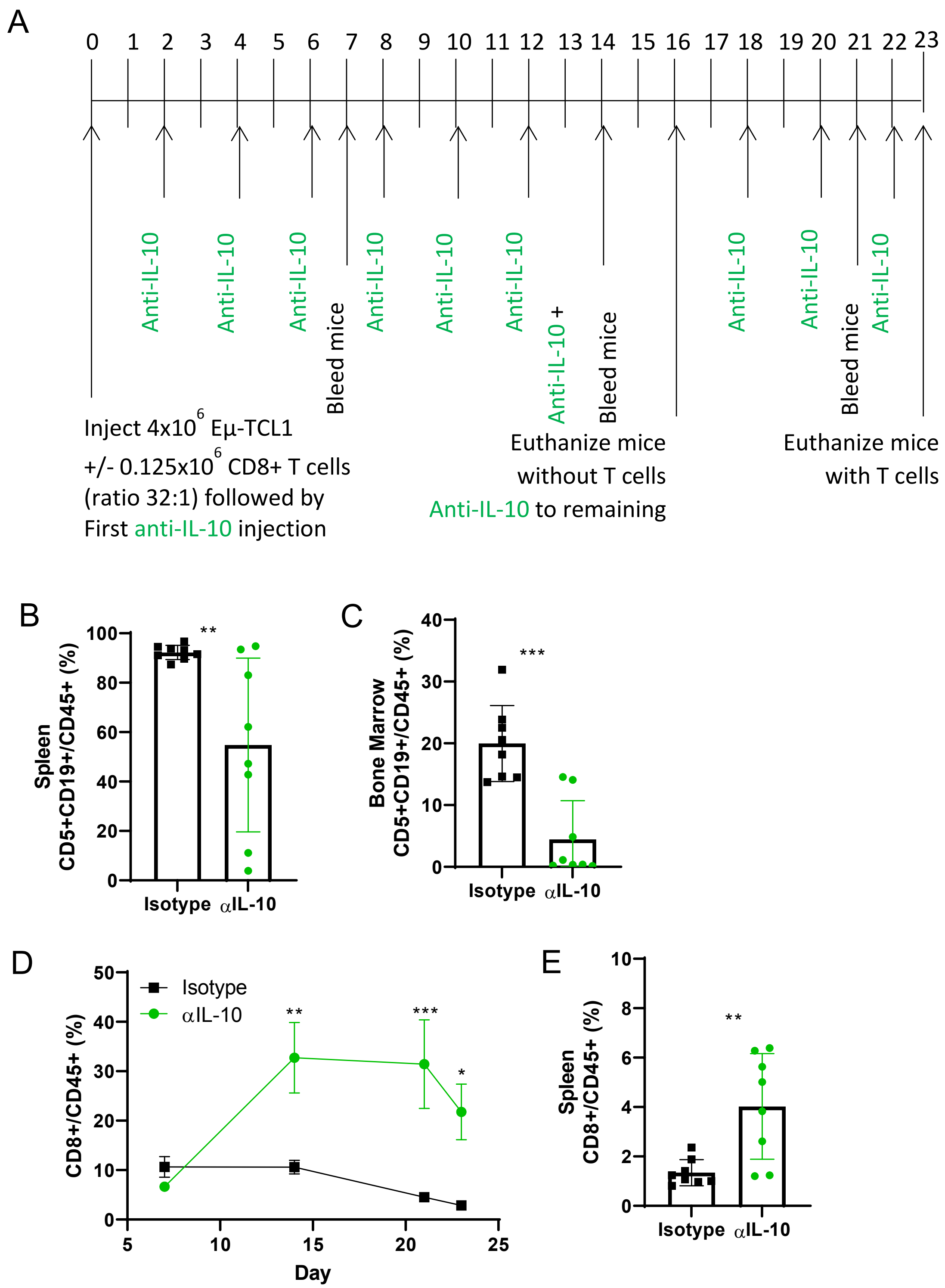

### Supplementary Figure 5

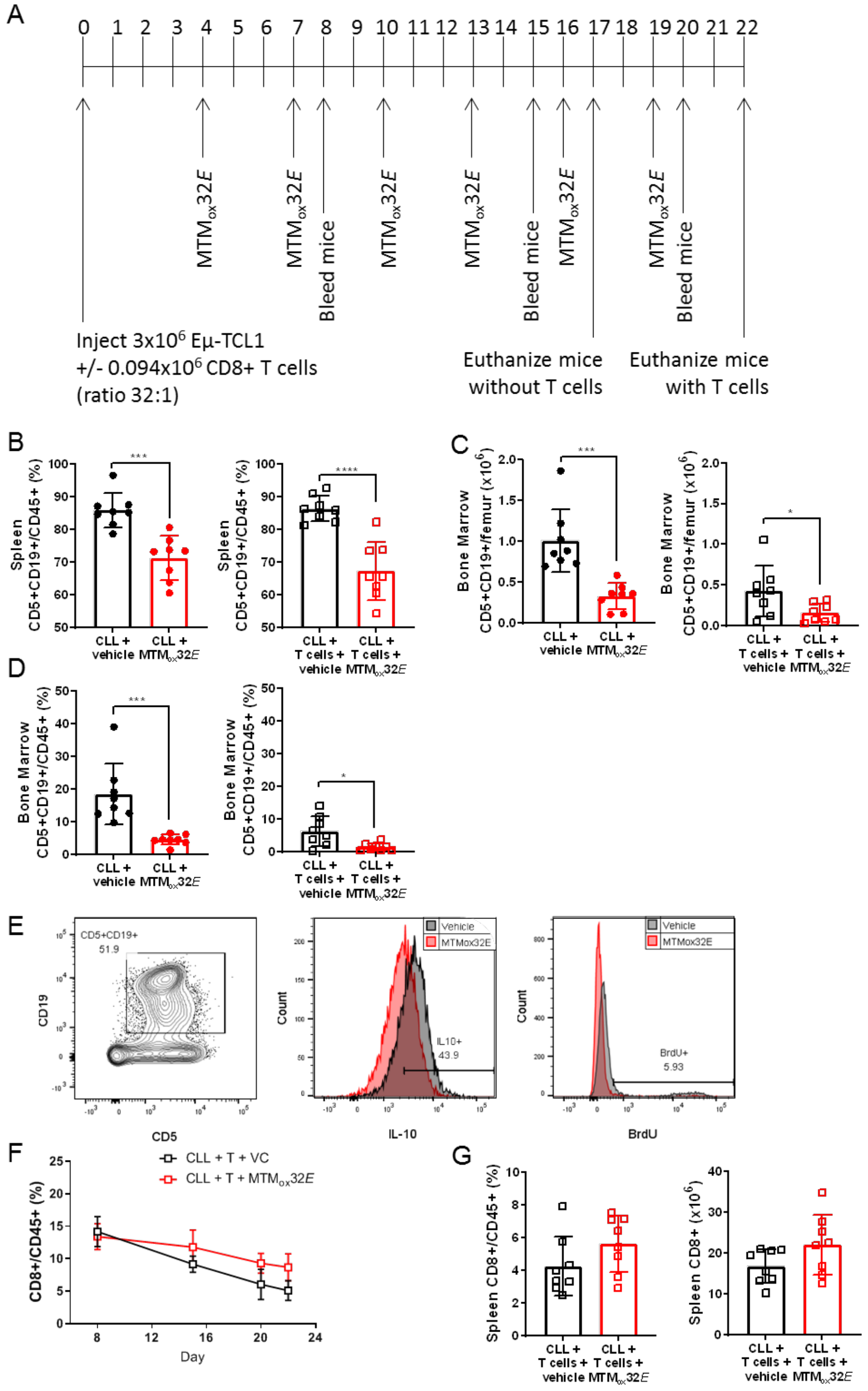

**Supplementary Figure 6**

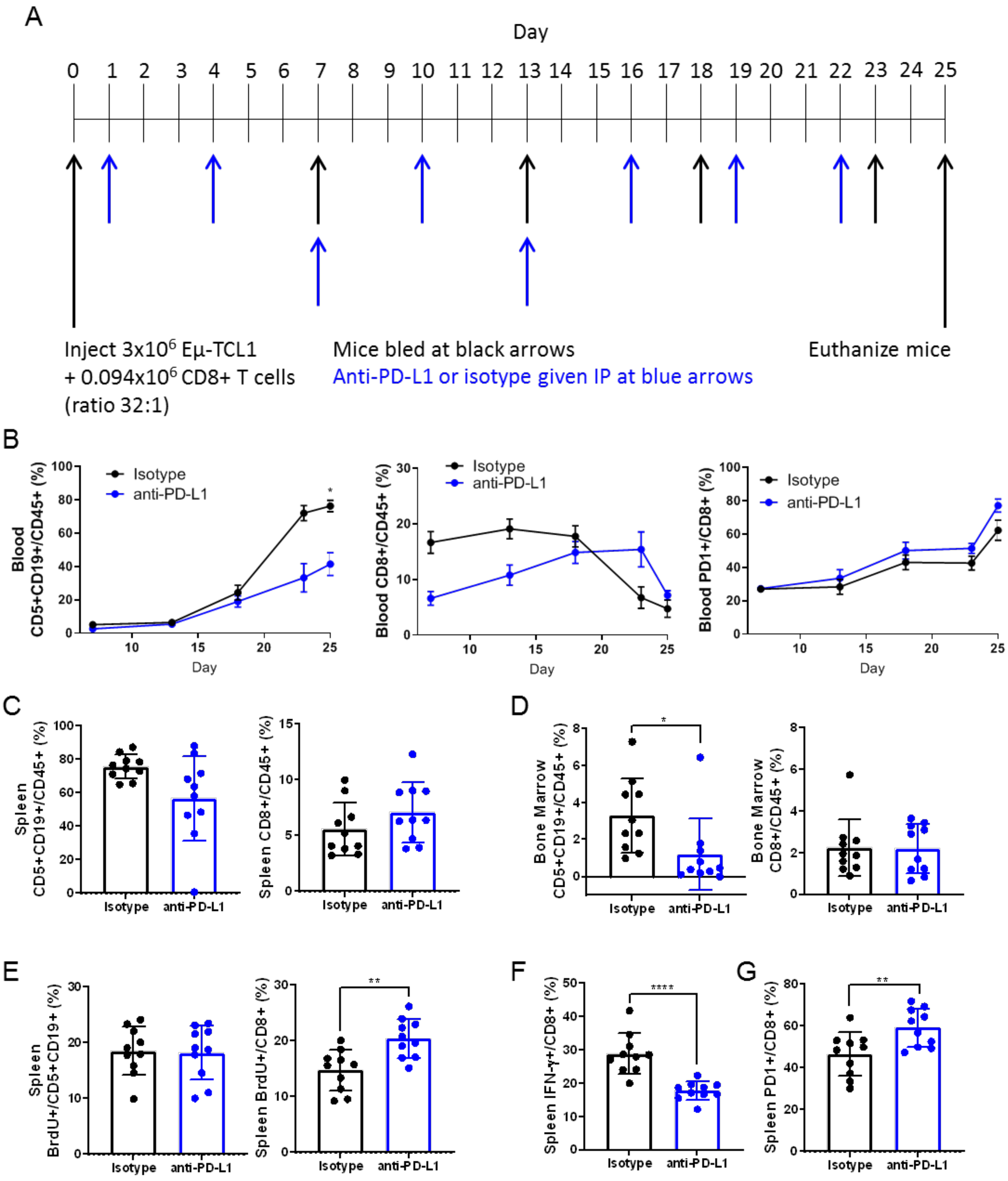

1 **Supplementary Figure 7**

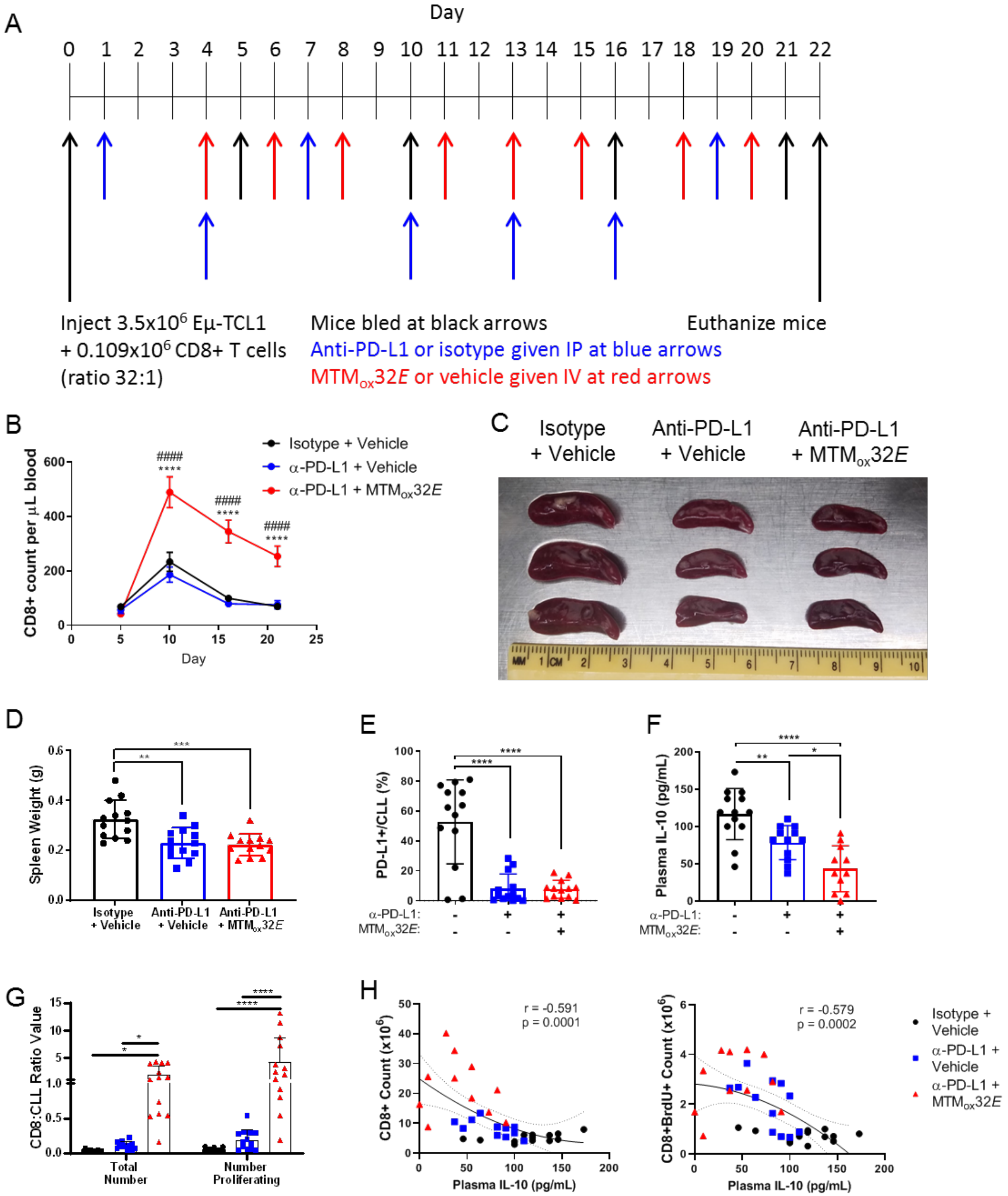


2

### Supplementary Figure 8

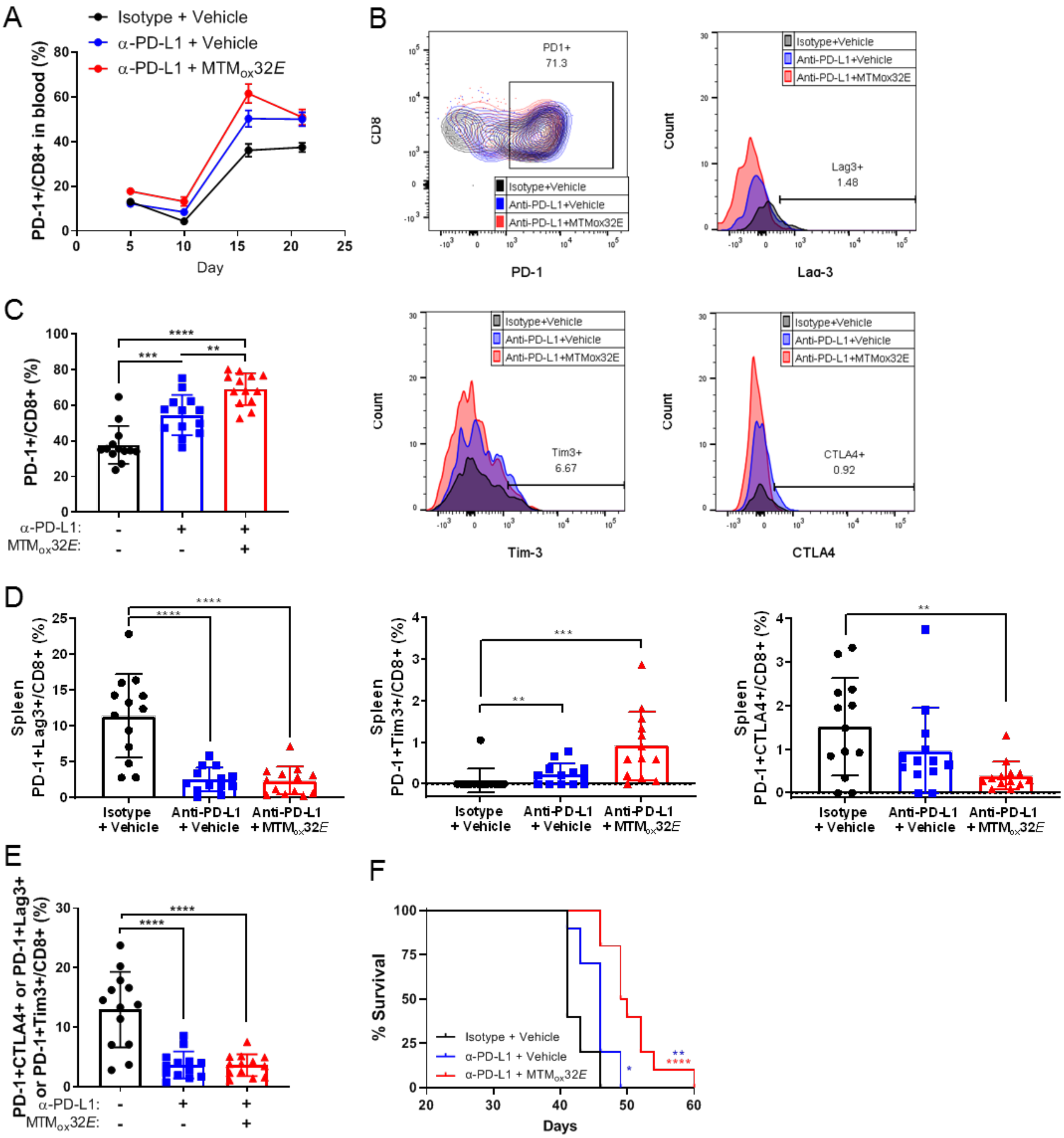

**Supplementary Table 1**

| Patient | Age | Race | Gender | Past Treatment | Current Treatment | WBC (k/uL) | CD5+19+ | CD38+ | Zap70+ | CD49d+ | IgH | Isotype |
|---|---|---|---|---|---|---|---|---|---|---|---|---|
| hCLL20 | 57 | W | M | None | None | 7.4 | 44.40% | ND | ND | ND | U-CLL | IgM |
| hCLL26 | 62 | W | M | None | None | 62.5 | 90.80% | No | Yes | No | M-CLL | IgM |
| hCLL28 | 58 | W | M | BR | None | 8.4 | 66.70% | No | Yes | No | U-CLL | IgM |
| hCLL29 | 80 | W | M | FCR | None | 39.6 | 91.70% | Yes | Yes | No | U-CLL | IgM |
| hCLL30 | 52 | W | M | None | None | 68.2 | 89.60% | No | Yes | No | U-CLL | IgM |
| hCLL32 | 82 | W | M | Chlorambucil, BR | IVIG | 5.6 | 52.30% | Yes | Yes | No | U-CLL | IgM |
| hCLL33 | 59 | W | M | None | None | 21.3 | 78.50% | No | No | No | M-CLL | ND |
| hCLL34 | 66 | W | F | None | None | 11.8 | 67.10% | No | No | No | M-CLL | IgG |
| hCLL37 | 63 | B | F | None | None | 130 | 89.70% | No | No | No | U-CLL | IgM |
| hCLL38 | 57 | W | M | None | None | 64.7 | 86.00% | No | No | No | U-CLL | IgM |
| hCLL39 | 75 | W | M | None | None | 22.3 | 76.30% | No | No | No | M-CLL | IgM |
| hCLL40 | 78 | W | F | None | None | 12.1 | 37.80% | No | No | No | M-CLL | IgM |
| hCLL42 | 79 | W | M | BR | None | 6.7 | 64.60% | No | No | No | M-CLL | IgM |
| hCLL43 | 68 | W | M | None | None | 16.8 | 75.80% | Yes | No | No | M-CLL | IgM |
| hCLL44 | 60 | W | F | None | None | 16.8 | 54.80% | No | No | No | U-CLL | IgM |
| hCLL47 | 71 | W | M | None | Ibrutinib-6 wk | 90.8 | 34.60% | Yes | No | No | M-CLL | ND |
| hCLL51 | 62 | W | M | None | None | 220 | 45.50% | Yes | Yes | No | M-CLL | ND |
| hCLL54 | 47 | B | F | None | None | 12.1 | 94.30% | No | Yes | No | ND | ND |
| hCLL60 | 55 | W | M | None | None | ND | 91.80% | No | No | No | ND | ND |

**ND: not determined**

**Supplementary Table 2**

| BioLegend α-Mouse Antibodies | | BioLegend α-Human Antibodies | |
|---|---|---|---|
| **Antigen-Fluorophore** | **Product Number** | **Antigen-Fluorophore** | **Product Number** |
| BrdU-FITC | 364104 | CD19-PerCP/Cy5.5 | 302230 |
| CD107a-FITC | 121606 | CD25-PE | 302606 |
| CD19-APC | 115512 | CD38-PE/Cy7 | 356608 |
| CD19-PE/Cy7 | 115520 | CD4-PEdazzle594 | 300548 |
| CD25-FITC | 102005 | CD45-APC | 368512 |
| CD27-APC | 124212 | CD45-APC/Cy7 | 304014 |
| CD28-FITC | 122008 | CD49d-APC/Cy7 | 304328 |
| CD4-PacBlue | 100428 | CD5-FITC | 364022 |
| CD4-PE/Cy7 | 100528 | CD69-PE/Cy7 | 310912 |
| CD45-BV510 | 103138 | CD8a-APC | 301049 |
| CD45-PacBlue | 103126 | IFNγ-PacBlue | 502522 |
| CD45-PE/Cy5 | 103110 | Zap70-PE | 313404 |
| CD45-PE/Cy7 | 103114 | LEAF Purified CD28 | 302934 |
| CD5-FITC | 100606 | LEAF Purified CD3 | 300314 |
| CD5-PE | 100608 | LEAF Purified CD40 | 313010 |
| CD5-PE/Cy7 | 100622 | LEAF Purified IL-10 | 501504 |
| CD8-APC/Cy7 | 100714 | **BioLegend α-Mouse Antibodies** | |
| CD8-FITC | 100706 | **Antigen-Fluorophore** | **Product Number** |
| CD8-PacBlue | 100725 | PD-1(CD279)-PE | 135206 |
| CD8-PE | 100708 | PD-1(CD279)-PerCP-Cy5.5 | 135208 |
| CD8-PE/Cy7 | 100722 | PD-L1(CD274)-PE | 124308 |
| CTLA-4(CD152)-APC | 106310 | Tim-3(CD366)-PE/Cy7 | 134010 |
| Granzyme B-PerCP-Cy5.5 | 372212 | TNFα-APC | 506308 |
| IFNγ-FITC | 505806 | LEAF Purified CD28 | 102112 |
| IL-10-PE | 505008 | LEAF Purified CD3ε | 100331 |
| KLRG1-PerCP/Cy5.5 | 138418 | LEAF Purified CD40 | 102810 |
| Lag-3(CD223)-PE | 125208 | LEAF Purified IL-10 | 505002 |

| Jackson ImmunoResearch α-Mouse Antibody | | Jackson ImmunoResearch α-Human Antibody | |
|---|---|---|---|
| **Antigen** | **Product Number** | **Antigen** | **Product Number** |
| IgM, μ chain | 115-006-020 | IgM, Fc5μ Fragment | 109-006-129 |

**Supplementary Table 3**

| IC50 (nM) | Viability | | IL-10 | |
|---|---|---|---|---|
| | MTM | $MTM_{ox}32E$ | MTM | $MTM_{ox}32E$ |
| Eμ-Tcl1 | 159 | ~ | 44 | 69 |
| hCLL20 | ~ | ~ | 25.8 | 80.2 |
| hCLL26 | 182.9 | ~ | 32.6 | 80.0 |
| hCLL28 | 108.1 | 4.07 | 26.2 | 1914 |
| hCLL32 | 103.3 | 306.5 | 32.6 | 45.9 |
| hCLL33 | 50.8 | ~ | 36.9 | 1120 |
| hCLL34 | 68.0 | 50.0 | 154.7 | 324.9 |
| hCLL37 | 173.7 | 198.6 | ~ | 40.66 |
| hCLL38 | 169.4 | 1.168 | 93.0 | 19.0 |
| hCLL39 | 169.6 | ~ | 66.7 | ~ |
| hCLL40 | 94.7 | ~ | 24.5 | 301.2 |
| hCLL42 | 105.8 | ~ | 20.8 | 6120 |
| hCLL43 | 98.1 | ~ | 29.4 | 909.2 |
| hCLL44 | ~ | ~ | 20.96 | 304.2 |
| hCLL47 | ~ | 40.87 | 21.27 | 213.8 |
| hCLL51 | 92.53 | ~ | 40 | 328.6 |

| IC50 (nM) | Mouse CD4 | | Mouse CD8 | | Splenocytes | |
|---|---|---|---|---|---|---|
| | MTM | $MTM_{ox}32E$ | MTM | $MTM_{ox}32E$ | MTM | $MTM_{ox}32E$ |
| IFN-γ | 117.5 | 183.1 | 33.02 | ~ | 299 | ~ |
| IL-2 | 357.7 | ~ | 706.5 | ~ | | |
| IL-10 | 86.79 | ~ | | | | |
| Viability | 67.52 | ~ | 101.3 | 468.2 | 183 | ~ |

**~ Not measurable**

**Supplementary Figure 1: IL-10 reduces human CD8+ T-cell activation and $MTM_{ox}32E$ inhibits CLL IL-10 production.** (A-B) CD69 expression of healthy donor (A) or CLL patient (B) human CD8+ T-cells in response to a 5-day stimulation with plate bound anti-CD3 with soluble anti-CD28 and varying amounts of exogenous recombinant human IL-10, as in Fig. 1. (C-D) Response of hCLL CD8+ T-cells stimulated with StemCell Human T-cell activation kit for 5 days with 1µM $MTM_{ox}32E$. Percent of IFN-γ+ (C) or BrdU+ (D) CD8+ T-cells normalized to vehicle treated controls. (E-F) Response of healthy donor CD8+ T-cells stimulated with anti-CD3/anti-CD28 Dynabeads for 5 days with 1µM $MTM_{ox}32E$. Percent of IFN-γ+ (E) or BrdU+ (F) CD8+ T-cells normalized to vehicle treated controls. (G) Secreted IL-10 from Mec1 cells treated with MTM or $MTM_{ox}32E$ for 24 hours, normalized to vehicle control (630pg/mL). (H) Fold change in Mec1 mRNA for Sp1 target genes when treated with 100nM MTM or $MTM_{ox}32E$. (I) Fold change in hCLL mRNA for Sp1 target genes when treated with 100nM or 1000nM MTM and $MTM_{ox}32E$. Panel shows a box and whisker plot showing mean, 25$^{th}$-75$^{th}$ percentile, and overlaid with dots representing average values for each CLL patient (n=5). (J-L) IL-10 secretion (J) or viability (K-L) of hCLL PBMCs (by resazurin fluorescence) cultured 24 hours with 25µg/mL soluble anti-IgM and different doses of MTM (J+L) or $MTM_{ox}32E$ (K), normalized to vehicle control. Statistical comparisons were performed: in A-G and I by two-way ANOVA for 0 ng/mL IL-10 to varying concentrations (A+B), vehicle to $MTM_{ox}32E$ (C-F), or MTM to $MTM_{ox}32E$ (G+I), and in H by one-way ANOVA for MTM to $MTM_{ox}32E$. *$p<0.05$, **$p<0.01$, ***$p<0.001$, ****$p<0.0001$

**Supplementary Figure 2: $MTM_{ox}32E$ and MTM have similar effects on Eµ-TCL1 IL-10 and viability but differing effects on murine $CD4^+$ and $CD8^+$ T-cells.** (A) IL-10 secretion from Eµ-TCL1 splenocytes when treated with MTM or $MTM_{ox}32E$ for 24 hours. (B) Anti-CD3 stimulated T-cell cytokine secretion ($CD4^+$ IL-2 or $CD8^+$ IFN-γ) when treated with MTM or $MTM_{ox}32E$ for 24 hours. (C) Viability of Eµ-TCL1 splenocytes (by resazurin fluorescence) when treated with MTM or $MTM_{ox}32E$ for 24 hours. (D) Anti-CD3 stimulated T-cell viability ($CD4^+$ or $CD8^+$) when treated with MTM or $MTM_{ox}32E$ for 24 hours. (E-F) Viability of anti-CD3 stimulated murine C57BL/6 CD4+ T-cells, CD8+ T-cells or unstimulated Eμ-TCL1 cells (by resazurin fluorescence) when treated with $MTM_{ox}32$*E* (E) or MTM (F) for 24 hours. Statistical comparisons made by two-way ANOVA between points on different lines at the same drug concentrations. *p<0.05, **p<0.01, ***p<0.001, ****p<0.0001

**Supplementary Figure 3: Concentrations of $MTM_{ox}32$*E* that inhibit CLL IL-10 production have only modest effects on human T-cell functionality.** (A) Viability of healthy human PBMCs as determined by Annexin V and PI staining after five days in culture with $MTM_{ox}32$*E*. (B) Representative histograms of cytokine production by healthy human $CD4^+$ (left) and $CD8^+$ (right) T-cells. (C) Quantification of cytokine production by human T-cells out of mixed PBMC cultures (top) or by sorted human CD4+ T-cells (bottom) after five days of treatment with $MTM_{ox}32$*E*. Statistical significance was calculated by one-way ANOVA for control (untreated) to $MTM_{ox}32$*E*. *p<0.05.

**Supplementary Figure 4: Blocking IL-10 with antibody slows the progression of Eμ-TCL1 CLL in NSG mice.** (A) Experiment timeline for Fig. 2E+2G. Antibody was administered IP at 150ug/mouse per injection. n=8 mice/group (B-C) Frequency of CLL cells in spleen (B) and bone marrow (C) of NSG mice given Eμ-TCL1 and primed $CD8^+$ T-cells. (D-E) Frequency of CD8+ T-cells as a function of time after CLL injection in the blood (D) and spleen (E) of NSG mice given Eμ-TCL1 and primed $CD8^+$ T-cells (mean + SEM in D). Statistical comparisons in B, C, E were obtained by unpaired student's T-test and in D by two-way ANOVA between isotype and anti-IL-10. *p<0.05, **p<0.01, ***p<0.001

**Supplementary Figure 5: Inhibiting CLL IL-10 with $MTM_{ox}32$*E* slows the progression of Eμ-TCL1 CLL in NSG mice.** (A) Experiment timeline for Fig. 3 and 4. (n=8 mice/group) (B) Frequency of Eμ-TCL1 CLL cells in the recipient spleen. (C-D) Count (C) and frequency (D) of Eμ-TCL1 CLL cells in the bone marrow. (E) Representative gating and histograms of IL-10+ or BrdU+ CLL cells from the spleen. (F) Frequency of $CD8^+$ T-cells in the blood (mean + SEM). (G) Frequency and number of $CD8^+$ T-cells in the spleen. Statistical significance was calculated by unpaired student's t-test in B-E and by two-way ANOVA for vehicle to $MTM_{ox}32E$ in F. *p<0.05, **p<0.01, ***p<0.001, ****p<0.0001

**Supplementary Figure 6: Checkpoint blockade alone slows the progress of Eμ-TCL1 CLL in NSG mice but is not a strong enhancer of T-effector cell function.** (A) Experiment timeline. (B) Frequency of Eμ-TCL1 CLL cells (left), $CD8^+$ T-cells (middle), and $PD\text{-}1^+$ $CD8^+$ T-cells (right) in the blood of NSG mice (mean + SEM). Graphs C-G show data from mice after euthanasia. (C) Frequency of Eμ-TCL1 CLL cells (left) and $CD8^+$ T-cells (right) in the spleen. (D) Frequency of Eμ-TCL1 CLL cells (left) and $CD8^+$ T-cells (right) in the bone marrow. (E) Frequency of $BrdU^+$ Eμ-TCL1 CLL cells (left) and $CD8^+$ T-cells (right) in the spleen. (F-G) Frequency of $IFN\text{-}\gamma^+$ (F) and $PD\text{-}1^+$ (G) $CD8^+$ T-cells in the spleen. Statistical comparisons in B were performed by two-way ANOVA for vehicle to $MTM_{ox}32E$ at each day sampled, and in C-G by unpaired student's t-test. *p<0.05, **p<0.01, ****p<0.0001

**Supplementary Figure 7: Checkpoint blockade is more effective when combined with IL-10 suppression.** (A) Experiment timeline for Fig. 5, 6 and 7. (n=13 mice/group). (B) Number of $CD8^+$ T-cells in the blood (mean + SEM). (C) Representative image of spleens from NSG mice. (D) Weight of whole spleens from NSG mice. (E) Frequency of $PD\text{-}L1^+$ Eμ-TCL1 CLL cells in the spleen of NSG mice. (F) Plasma IL-10 levels at euthanasia. (G) Ratio of the number of proliferating $CD8^+$ T-cells to the number of Eμ-TCL1 CLL cells. (H) Correlation between plasma IL-10 levels and total $CD8^+$ T-cell count (left) or proliferating $CD8^+$ T-cell count (right). Graphs show the Pearson's correlation coefficient, two-tailed p-value, trendline, and 95% confidence interval. Statistical comparisons in B were performed by two-way ANOVA for isotype + vehicle to either treatment group at each day sampled, and in D-G by one-way ANOVA for isotype + vehicle to either treatment group. *p<0.05, **p<0.01, ***p<0.001, and ****p<0.0001 indicate statistical significance (*difference between combination and ICB alone, #difference between combination and control).

**Supplementary Figure 8: CD8$^+$ T-cells are more functional when the NSG recipients are treated with both checkpoint blockade and MTM$_{ox}$32*E*.** (A) Frequency of PD-1$^+$ cells out of CD8$^+$ T-cells in the blood (mean + SEM). (B) Representative histograms showing reduced expression of Lag3, Tim3 and CTLA4 on whole CD8$^+$ T-cells in the group with combination treatment. (C) Frequency of PD-1$^+$ CD8$^+$ T-cells in the spleen. (D) Frequency of exhausted T-cells in the spleen, either PD-1$^+$Lag3$^+$ (left), PD-1$^+$Tim3$^+$ (middle), or PD-1$^+$CTLA4$^+$ (right). (E) Combined frequency of CD8$^+$ T-cells expressing PD-1 and an additional exhaustion marker. (F) Long term survival of mice with combination or monotherapy. NSG mice were given 4x10$^6$ CLL plus primed CD8$^+$ T-cells at 32:1 CLL:T-cells on day 0 (n=10/group). 10mg/kg anti-PD-L1 or isotype control was given IP every 3 days from day 1 to day 22. 12mg/kg MTM$_{ox}$32*E* or vehicle (5% Kolliphor 1% DMSO) was given IV every 2-3 days from day 4 to day 22. CLL burden was monitored in the blood and after mice reached 75% CD5+CD19+ CLL, body condition was monitored, and mice were euthanized at a body condition score of 2 or greater. Significance: blue stars by blue line: isotype+vehicle vs anti-PD-L1+vehicle; blue stars by red line: isotype+vehicle vs anti-PD-L1+MTM$_{ox}$32*E*; red stars by red line: anti-PD-L1+vehicle vs anti-PD-L1+MTM$_{ox}$32*E*. Statistical comparisons were performed by two-way ANOVA for isotype+vehicle to either treatment group in A, one-way ANOVA for isotype+vehicle to either treatment group in C-E, and Log-rank Mantel-Cox test in F. *p<0.05, **p<0.01, ***p<0.001, ****p<0.0001

**Supplementary Table 1: Human CLL patient data.** Total white blood cell (WBC) counts and the percent of CD5+CD19+ cells at the time of collection are indicated. Threshold for CD38, Zap70, and CD49d positivity was set at 30% of CD5$^+$CD19$^+$ events. Abbreviations: BR = Bendamustine/Rituxan, FCR = Fludarabine/Cyclophosphamide/Rituximab, IVIG = Intravenous Immunoglobulin, U- = unmutated BCR, M- = mutated BCR, ND = not determined

**Supplementary Table 2: Antibodies used in experiments.** Antibodies used are organized by company and listed with product numbers. Legend: LEAF = low endotoxin azide free, PacBlue = Pacific Blue

**Supplementary Table 3: IC50 of $MTM_{ox}$32*E* on cell culture viability and cytokine secretion.** Blank spaces were left if the test was not performed. ~ means IC50 could not be calculated in the dose range tested. IC50 values were calculated using GraphPad Prism 8 software.
